# The complex genetic architecture of recombination and structural variation in wheat uncovered using a large 8-founder MAGIC population

**DOI:** 10.1101/594317

**Authors:** Rohan Shah, B Emma Huang, Alex Whan, Marcus Newberry, Klara Verbyla, Matthew K Morell, Colin R Cavanagh

## Abstract

**Background:** Identifying the genetic architecture of complex traits requires access to populations with sufficient genetic diversity and recombination. Multi-parent Advanced Generation InterCross (MAGIC) populations are a powerful resource due to their balanced population structure, allelic diversity and enhanced recombination. However, implementing a MAGIC population in complex polyploids such as wheat is not trivial, as wheat harbours many introgressions, inversions and other genetic factors that interfere with linkage mapping.

**Results:** By utilising a comprehensive crossing strategy, additional rounds of mixing and novel genotype calling approaches, we developed a bread wheat eight parent MAGIC population made up of more than 3000 fully genotyped recombinant inbred lines derived from 2151 distinct crosses, and achieved a dense genetic map covering the complete genome. Further rounds of inter-crossing led to increased recombination in inbred lines, as expected. The comprehensive and novel approaches taken in the development and analysis of this population provide a platform for genetic discovery in bread wheat. We identify previously unreported structural variation highlighted by segregation distortion, along with the identification of epistatic allelic interactions between specific founders. We demonstrate the ability to conduct high resolution QTL mapping using the number of recombination events as a trait, and identify several significant QTLs explaining greater than 50% of the variance.

**Conclusions:** We report on a novel and effective resource for genomic and trait exploration in hexaploid wheat, that can be used to detect small genetic effects and epistatic interactions due to the high level of recombination and large number of lines. The interactions and genetic effects identified provide a basis for ongoing research to understand the basis of allelic frequencies across the genome, particularly where economically important loci are involved.

## Background

Due to the large genome size of bread wheat and its hexaploid nature, powerful genetic resources are required to understand the underlying genetic mechanisms for a wide variety of phenotypes. While a variety of biparental and association populations have been developed for this purpose, multiparental populations offer a novel opportunity to dissect genomic structure by combining strengths of both prior approaches. In particular, Multiparent Advanced Generation Inter-Crosses (MAGIC) mix the genomes of several diverse founders through multiple generations of intercrossing and selfing or double haploidy, to generate a large population of immortalised lines.

MAGIC populations have been developed as genetic resource panels in a number of species [Cavanagh et al., 2008, Huang et al., 2015], but have seen the greatest uptake in wheat in terms of both number of different populations and size thereof. The first plant MAGIC population was developed in Arabidopsis thaliana [Kover et al., 2009], and since then populations have been developed in crops including barley [Mathew et al., 2018], chickpea [Gaur et al., 2012], rice [Bandillo et al., 2013], ryegrass [Begheyn et al., 2018] and tomato [Pascual et al., 2015]. None of these populations in other crops explore the full range of potential intercrosses possible in the early stages of MAGIC designs, and besides [Bandillo et al., 2013] none of them have been genotyped on more than 1000 lines. By contrast, the four-parent spring wheat MAGIC population [Huang et al., 2012] consists of nearly 1500 RILs genotyped at high density, while the eight-parent winter wheat MAGIC [Mackay et al., 2014] consists of 1091 F7RILs, of which 720 have been genotyped with the 90K SNP chip [Wang et al., 2014]. This wealth of genetic data and genetic diversity facilitates the use of these populations for uncovering genomic structures.

Thus far, MAGIC populations have been analyzed primarily with a focus on linkage map construction and QTL mapping. High-density linkage maps have been constructed in wheat [Huang et al., 2012, Cavanagh et al., 2013, Gardner et al., 2016, Sannemann et al., 2018], durum wheat [Milner et al., 2016], barley [Sannemann et al., 2015] and tomato [Pascual et al., 2015], and validated against consensus and physical maps. The diversity and resolution of MAGIC populations enables the mapping of more markers, more precisely, than in previous populations. The resulting maps provide greater resolution in QTL mapping, which has been performed in all of the crops previously mentioned, both for proof of concept [Kover et al., 2009, Huang et al., 2012, Sannemann et al., 2015] and discovery of novel loci [Rebetzke et al., 2014, Barrero et al., 2015].

The rich genomic information contained in these populations enables investigation of genomic structure at a level of detail not possible in biparental populations. Multi-parental populations have been used to demonstrate widespread genetic incompatibilities in Drosophila, Arabidopsis, and maize [Corbett-Detig et al., 2013], and characterize regions associated with maternal transmission ratio distortion in mice [Didion et al., 2015]. In wheat, they have previously been used to identify widespread segregation distortion and introgressions [Gardner et al., 2016].

The production of a high-quality reference sequence for bread wheat has been a challenging problem, due to its large genome size and three separate subgenomes. The recent publication of the first reference sequence [The International Wheat Genome Sequencing Consortium (IWGSC), 2018] has underlied more recent work on the bread wheat genome [Keeble-Gagnère et al., 2018, Ramírez-González et al., 2018]. However, genetic maps constructed from large populations remain useful for the investigation of non-mendelian inheritance and for highlighting regions of uncertainty in the sequence data [Deokar et al., 2014].

In this paper we present an eight parent spring bread wheat MAGIC population. Three founders (Baxter, Westonia, and Yitpi) are Australian cultivars, which were previously used in a four-parent MAGIC population [Huang et al., 2012]. The other five founders originated from Canada (AC Barrie); USA (Alsen); CIMMYT (Pastor); Israel (Volcani); and China (Xiaoyan54). All founders are spring wheats with the exception of Xiaoyan54 which is a winter wheat. Founders were chosen on the basis of genetic and phenotypic diversity, with a particular emphasis on diversity for wheat quality traits. All except Volcani have been grown commercially.

Our single population contains three subpopulations, which have been produced with different levels of intercrossing in the mixing phase, allowing the assessment of the advantages such an approach gives. All of these subpopulations are used for linkage map construction, resulting in a dense and high-resolution map, which can be used to link genetic and physical maps for wheat. The completeness of the funnel structure and size of the population provide power to dissect the genetic structure of wheat, including genetic interactions and localized segregation distortion.

## Results and discussion

### Genotype calling

Genotyping was performed for lines from all populations using the Infinium iSelect 90K [Wang et al., 2014] SNP assay. This resulted in data from 81,587 markers. At the end of the marker calling process there were 29,566 polymorphic markers, of which 6,743 were reviewed manually. Figure S1 shows the number of markers called using each marker calling strategy. Method “DBSCAN-n” indicates the use of DBSCAN, where *n* is the number of alleles identified by DBSCAN.

For the markers called using HBC, the proportion of *marker heterozygote* calls is shown in Figure S2. The theoretically expected proportion of *identity by descent heterozygotes* for five generations of selfing is 0.0312 without intercrossing, and 0.0273 with intercrossing. Note that *marker heterozygotes* are not the same as identity-by-descent heterozygotes; if two parents carry the same marker allele, then a heterozygote for those parents will not be a marker heterozygote. So the proportion of marker heterozygotes is lower than the proportion of identity-by-descent heterozygotes, and Figure S2 shows a situation where the proportion of marker heterozygotes is generally higher than expected. This is partially due to our fairly aggresive calling of marker heterozygotes. The higher rate of heterozygote calls could also be due to the presence of homologues. A proper assesement of heterozygosity will be made by using identity-by-descent hetoryzogetes, using all markers *jointly*, in subsequent analysis.

Calling for most markers was straightforward, however there were a small number of challenging cases that were initially miscalled. These cases are only identifiable *after* a genetic map has been constructed. For example, marker Kukri c37840 253 is present on chromosomes 2A and 2B, and polymorphic on both, although with a genetic map this is not obvious (Figure S3a). In Figure S3b, color represents the imputed genotype at a location on chromosome 2B. In Figure S3c, color represents the imputed genotype at a location on chromosome 2A. As this pattern cannot reasonably have arisen by chance, it is clear that there are four clusters here. Without a genetic map there will appear to be only two clusters, and calling of this marker will be incorrect. The marker will appear to map to two chromosomes, although due to the incorrect calling it will not be correct on either chromosome.

Table S1 summarises the number of markers with segregation distortion, on each chromosome. However, this potentially says more about the markers than the underlying genetic structure; the ability to successfully call marker alleles varies across markers, and across marker alleles for each marker. So observed segregation distortion at the individual marker level is susceptible to effects relating to the marker calling algorithm, and the ease with which different marker alleles can be called. A better investigation of *genetic* distortion will be made using identity-by-descent probabilities, using all markers *jointly*, in subsequent analysis.

### Map summary

Our constructed map has 27,687 markers across all 21 chromosomes. Table S1 provides a complete summary of the mapped markers. Markers that are distorted are still present in the map. Removing markers on the basis of single-locus distortion is difficult, as the distortion may be due only to a difference in the marker calling rate for different marker alleles.

Figure S4 shows the positions of all markers, and all gaps in the map, with color representing the number of unique positions per centiMorgan. Figure S5 is similar, but color represents the number of markers per centiMorgan. Chromosome 2B has been estimated as being very long; this is an unavoidable result of the large number of markers on that chromosome. As marker density increases, distances between adjacent markers become extremely small. As a result, any estimation error will almost certainly be in the direction of estimating a distance that is too large. Compounding the problem, there are more such estimates to be made. Genotyping error may also lead to the separation of markers that do not have any true recombination event separating them. We note that the relevant inputs to QTL mapping algorithms, such as genotype probabilities and imputed genotypes, are relatively insensitive to overall chromosome length.

The marker densities of 1.58 unique positions / cM on the A genome and 2.02 unique positions / cM on the B genome were far higher than the marker density of 0.50 unique positions / cM for the D genome. Around 37%, 55% and 6% of markers were mapped to the A, B and D genomes respectively. The lower coverage on the D chromosomes is reflected both in the length (parts missing) and the resolution (bigger gaps). This feature is known to reflect the lower polymorphism of the D genome [Wang et al., 2014]. However, it *may* also be related to the presence of markers polymorphic on multiple chormosomes. For example, we have noted a large number of markers polymorphic on both 2B and 2D. These markers are not simple to map, and have not been included. If a large fraction of markers on the D genomes are also present on the A or B genomes, and cannot be mapped, this could be interpreted as reduced polymorphism on the D genome.

In general, the number of unique positions per chromosome was far lower than the number of markers mapped. The 27,687 markers were mapped to 7,674 unique positions.

### Map comparison

The map constructed in this population has the highest combination of coverage, density and resolution of any constructed from a single population in wheat. While the 90K consensus map contains nearly 47,000 markers, each of the eight individual biparental populations contributing to this map contains fewer than 19,000 markers. A previously published consensus map [Li et al., 2015] has 28,000 markers at 3,757 unique positions, but for each of the three populations merged to form the consensus, fewer than 20,000 markers were mapped. Genotyping by sequencing can detect a very high number of markers, resulting in genetic maps with extremely high marker density. In wheat, GBS-based maps have been reported with nearly 20,000 SNPs in 1,485 bins [Poland et al., 2012], over 400,000 in 1421 bins [Saintenac et al., 2013], and 1.7 million markers in 1335 bins [Chapman et al., 2015]. In the last example, the map was constructed from a doubled haploid population of 90 lines, demonstrating that the number of unique locations is limited by the recombination observed in the population. One of the significant advantages of MAGIC populations is the high number of recombination events due to the multiple rounds of crossing. A previously reported map developed with 643 lines from an 8-way wheat MAGIC population (referred to subsequently as NIAB 8-way Gardner et al. [2016]) has 18,601 markers in 4,578 unique positions.

### Map validation

Figure S6 shows the comparison of the MAGIC map with the consensus map based on the 90K SNP chip [Wang et al., 2014]. Points in blue represent markers that are within 20cM vertically of the line of best fit. Other (conflicting) markers are shown in red. Figure S7 shows a similar figure for the NIAB 8-way.

A summary of the markers with agreeing and conflicting positions can be found in Table S2. The comparisons show a large number of conflicts on chromosome 2B. This is due to the Sr36 introgression [Tsilo et al., 2008]. The introgression is rarely broken up by recombination events (see discussion of recombination), and displays distorted inheritance (see discussion of segregation distortion main effects). This makes map construction for chromosome 2B difficult using this population.

Figure S8 compares our map and the IWGSC RefSeq v1.0. Large disagreements between the physical map and the MAGIC map may indicate differences in genome ordering between Chinese Spring and the parents used for mapping. They may also be due to paralogous sequences, or genetic insertions, deletions and inversions. On a fine scale, the order of markers in the genetic map and the IWGSC RefSeq v1.0 are not expected to match exactly, due to variability in recombination fraction estimates affecting the ordering in the MAGIC map. We do not expect to have high enough resolution in the MAGIC map to match the resolution of the sequence data; it has previously been demonstrated that a genetic map is insufficient to completely order scaffolds in tomato [Shearer et al., 2014]. However, the high resolution of the MAGIC map can be used to highlight regions of uncertainty or dissonance in the physical map and improve reference assemblies [Deokar et al., 2014].

### Recombination

Comparison of the three subpopulations allows assessment of the benefit of the additional intercrossing, since AIC2RIL and AIC3RIL have two and three extra rounds of crossing, respectively. Table S3 shows the average number of recombination events for each chromosome, for each of the three subpopulations. Additional generations of intercrossing lead to a noticeable increase in recombination events. The imputed number of recombination events for chromosome 3B was much higher than the value of 2.6 in [Choulet et al., 2014], highlighting the high level of recombination in the MAGIC population. Figure S9 shows the distribution of the number of recombination events, for all subpopulations and the entire population.

The imputed haplotype blocks were used to estimate the average haplotype block size, at different points on the genome, for all three subpopulations. The haplotype block sizes were represented as a proportion of the chromosome size, and the results for the A and B genomes are shown in Figure S10. The decrease in block sizes toward either end of the chromosome is an obvious edge effect; the sizes of haplotype blocks near the ends of the chromosomes are limited by the edges of the chromosome. The two chromosomes with the largest average haplotype block size, as a proportion of chromosome length, are chromosomes 2B and 6B; both these chromosomes contain introgressions that distort genetic inheritance.

The haplotype blocks on chromosome 2B are particularly interesting. Figure S11 shows the average sizes of the haplotype blocks, classified according to their underlying genotype. The influence of the Sr36 introgression is obvious in the much larger size of the Baxter haplotype blocks. Figure S11 also shows an interesting spike in the average size of the Volcani haplotype blocks around 400 cM. This is difficult to interpret, as the difference seems to be due to a *decrease* in the number of small Volcani haplotype blocks at around 400 cM.

The results of performing QTL mapping, using the number of recombination events as a trait, are shown in Tables S4 - S7. For the number of recombination events on the B subgenome, the number of recombinations on chromosomes 2B and 6B was excluded, as these chromosomes are known to have large introgressions. QTL mapping was performed with these four traits, using the whole-genome approach [Verbyla et al., 2014]. We report only those QTL which explain over 1% of the phenotypic variance.

We found QTL explaining a total of 50.28%, 73.05% and 20.27% of variance for the A, B and D sub-genomes, respectively, and 37.37% for the whole genome. For the A subgenome, positions on the first half of chromosome 1A are highly significant, as are positions at the start of chromosome 3A. For the B subgenome, we find extremely large effects on chromosomes 5B and 7B. For the D genome, there are significant positions on chromosomes 3D and 7D. For the genome as a whole, we identify QTL on chromosomes 2A, 2B, 5B and 7B; recall that recombination on chromosome 2B was not counted as part of the B-subgenome recombination trait, but *is* included in the number of recombination events across the whole genome.

There is potential for very significant confounding effects when analysing recombination as a trait. The trait is a function of the imputed genotypes, as the trait is a count of the number of times the imputed genotype changes. If such confounding is present, we would expect to detect QTL that are artifacts of this functional relationship, and would have no meaning in terms of the underlying genetics. These would be likely to appear as QTL on a particular subgenome associated with variance for recombination events on the same subgenome. To check this, we simulated genetic data according to our pedigree and genetic map. We then counted recombination events for each line from the simulated data, and performed QTL mapping using the simulated trait and genetic data. We found that all detected QTL from simulated data explained less than 1% of the phenotypic variance, and also found no tendency for QTL for specific subgenome recombination traits to be on the same subgenome. This suggests that confounding effects are minimal in our analysis.

Another potential source of error in determining the number of recombinations could be incorrect marker ordering. If the marker ordering were wrong, we might see the imputed underlying genotype switch repeatedly between two genotypes. By contrast, in the correct ordering, the imputed underlying genotype might only change once. This badly ordered genetic region might then be detected as a QTL for recombination.

We do not believe that these errors contribute meaningfully to our results. Our genotype imputation method uses an error model, which can consider such repeated changes to be less likely than genotyping error, especially if the repeated changes happen in a small genetic region. Also, the artificially increased recombination would only occur within lines which actually had a recombination event within the badly ordered region. The number of such lines will be small, limiting the effect, because in regions with a large number of recombination events, genetic markers are easier to order. For these reasons, marker ordering errors are unlikely to reach high significance, especially for the wholegenome trait. By contrast, we find very highly significant effects; the QTL on chromosome 2B explain over 12% of genome-wide variation in recombination, and the QTL on chromosome 7B explain over 5% of genome-wide variation in recombination.

Analysis of a nested association mapping (NAM) population of 1,983 lines has previously found that variation in recombination is explained by a large number of rare alleles with small effects [Jordan et al., 2018]. In that NAM population, the average effect size was around 6.5% of variance explained, regardless of which trait was used. Accounting for the fact that our effects are estimated as spread over a number of genetic intervals, the QTL we report here are somewhat larger. We note that chromosome 7B was found to have the most QTL in the NAM population, with a region of 50 - 60 cM being identified as important in three separate biparental families. Excluding the 2B introgression, we also found that 7B had QTL with the largest effect, in approximately the same genetic region.

One possible reason for the larger effect sizes in our MAGIC population is the much higher genetic complexity; in our population all genetic variability is incorporated in a single population, whereas the NAM population is a collection of individual populations. Ultimately, this makes it difficult to draw direct comparisons.

### Segregation distortion

#### Main effects

Individual markers displaying distortion of segregation from that expected under Mendelian assumptions may indicate genotyping error. However, large groups of distorted markers may indicate biologically relevant phenomena, such as introgressions or translocations of genetic material. We have previously demonstrated that such regions can be mapped successfully in MAGIC populations [Huang et al., 2012, Shah et al., 2014].

Figure S12 gives a plot of the chi-squared statistics, testing for segregation distortion at the identity-by-descent level. We note (Table S1, Figure 1) a large region of segregation distortion on Chr 2B. At the most distorted point on chromosome 2B, Baxter is inherited by 26% of the final population, instead of the expected 12.5%. At the most distorted point, the chi-squared test statistic for this effect is 705, and the associated P-value is numerically equal to 0. In light of this P-value, and the presence of distortion across a large region of chromosome 2B, it is clear that this is real genetic effect. This distortion identifies an introgression known as Sr36 [Tsilo et al., 2008], which is contributed by the parent Baxter and is known to undergo meiotic drive.

**Figure 1:**
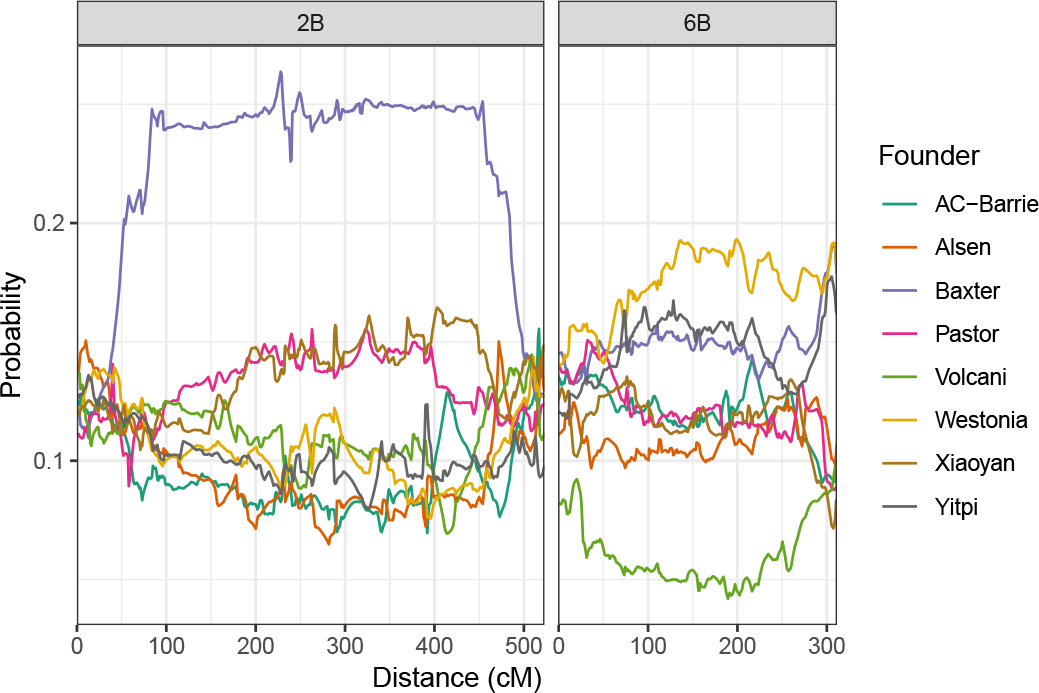
Average genetic composition across the entire population, for chromosomes 2B (left) and 6B (right).

Based on cytogentic analysis of Baxter (personal communications Cavanagh) the long arm of Chromosome 2B is replaced with 2G for Triticum Timopheevi. [Badaeva et al., 1996] showed that Chromosome 2G substituted for 2B at a frequency higher than expected, and suggested it may carry putative homoeoalleles of gametocidal genes present on group-2 chromosomes of several alien species. Chromosome 2G is recovered at a higher than expected frequency in the progeny of a hexaploid derivative of a cross between wheat and T. araraticum that was heterozygous for chromosomes 2G and 2B [Brown-Guedira et al., 1996].

The markers on chromosome 2B with the highest distortion are located between 299 cM and 322 cM in our map. This region was identified by first manually choosing ten markers known to be part of the introgression. These markers were all specific for the Baxter allele, and highly distorted. We then identified other markers specific for the Baxter allele, which were extremely strongly linked to the initial ten. As shown in Figures S4 and S5, this region contains a large number of markers.

The proportion of lines containing the introgression appears to depend on the number of generations of intercrossing (Table 1). It is clear that lines with intercrossing carry the introgression more frequently than those without intercrossing. It would also be expected that AIC3RIL lines should carry the introgression more frequently than AIC2RIL lines, but this is not observed; this may be due to the smaller sample size for the AIC2RIL and AIC3RIL subpopulations, compared to the MP8RIL subpopulation.

**Table 1:** Genetic composition at position 311 cM on chromosome 2B, which has the highest rate of Baxter alleles on chromosome 2B.

| Subpopulation | AC-Barrie | Alsen | Baxter | Pastor | Volcani | Westonia | Xiaoyan | Yitpi |
| --- | --- | --- | --- | --- | --- | --- | --- | --- |
| All | 0.08 | 0.09 | 0.26 | 0.14 | 0.11 | 0.09 | 0.14 | 0.09 |
| MP8RIL | 0.09 | 0.09 | 0.23 | 0.13 | 0.12 | 0.10 | 0.14 | 0.10 |
| AIC2RIL | 0.07 | 0.08 | 0.30 | 0.16 | 0.07 | 0.09 | 0.14 | 0.10 |
| AIC3RIL | 0.07 | 0.07 | 0.27 | 0.16 | 0.08 | 0.10 | 0.16 | 0.07 |

We also looked at the proportion of AIC2RIL lines carrying the introgression, out of those AIC2RIL lines for which a related AIC3RIL line carried the introgression (Table 2a). These AIC2RIL lines carry a Baxter allele at a much higher rate than on other non-distorted chromosomes. Similarly for those AIC2RIL lines *without* a related AIC3RIL line carrying the introgression (Table 2b).

**Table 2:** Genotypes of subsets of AIC2 lines, at several positions.

|  | Baxter | Other |
| --- | --- | --- |
| Sr36 | 0.39 | 0.61 |
| 1B, 150 cM | 0.27 | 0.73 |
| 1A, 150 cM | 0.31 | 0.69 |
| 5B, 150 cM | 0.22 | 0.78 |

|  | Baxter | Other |
| --- | --- | --- |
| Sr36 | 0.20 | 0.80 |
| 1B, 150 cM | 0.13 | 0.87 |
| 1A, 150 cM | 0.12 | 0.88 |
| 5B, 150 cM | 0.12 | 0.88 |

(b) Genetic composition of AIC2 lines at a position, for which a related AIC3 line has a non-Baxter genotype at the same position.

The presence of the introgression makes estimated recombination fractions somewhat unreliable, despite our use of a correction. As the estimation of map distances is based on estimated recombination fractions, this has the effect of inflating the length of chromosome 2B. Without a reliable model of genetic inheritance, there is little that can be done to fix this, short of rescaling the positions of all markers on that chromosome. The high marker density also contributes to the inflated length of this chromosome.

On chromosome 6B, Volcani is under-represented across most of the chromosome, being present in 4.2% of the final lines at position 190 cM, which is the point on 6B with the lowest rate of Volcani alleles. Westonia alleles are also inherited more frequently than expected, at this point. Table 3 shows the genetic composition of the population at 190 cM on chromosome 6B, for the entire population and all three subpopulations. The proportion of Volcani alleles decreases as the number of generations of intercrossing increases, with only 2% of AIC3RIL lines carrying the Volcani allele. This suggests that the Volcani allele is inherited less than 50% of the time in every generation.

**Table 3:**
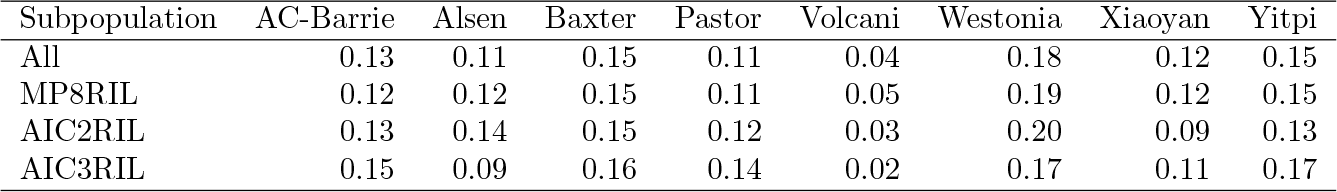
Genetic composition at position 190 cM on chromosome 6B, which has the lowest rate of Volcani alleles on chromosome 6B.

| Subpopulation | AC-Barrie | Alsen | Baxter | Pastor | Volcani | Westonia | Xiaoyan | Yitpi |
| --- | --- | --- | --- | --- | --- | --- | --- | --- |
| All | 0.13 | 0.11 | 0.15 | 0.11 | 0.04 | 0.18 | 0.12 | 0.15 |
| MP8RIL | 0.12 | 0.12 | 0.15 | 0.11 | 0.05 | 0.19 | 0.12 | 0.15 |
| AIC2RIL | 0.13 | 0.14 | 0.15 | 0.12 | 0.03 | 0.20 | 0.09 | 0.13 |
| AIC3RIL | 0.15 | 0.09 | 0.16 | 0.14 | 0.02 | 0.17 | 0.11 | 0.17 |

The chi-squared test statistic for distortion at 190 cM on chromosome 6B is 325, and the associated P-value is numerically equal to 0. In light of this P-value, and the presence of distortion across a large region of chromosome 6B, it is clear that this effect is statistically significant. As the distortion occurs almost chromosome-wide, it is not possible to determine the position of any genetic cause.

There is a further distortion on 6B, specific to the end of the chromosome. We estimate the position of this distortion as 308 cM. At this position, the Baxter, Westonia and Yitpi alleles are all inherited more frequently than expected. Table 4 gives the genetic composition at this point, for the entire population and all three subpopulations. Interestingly, additional generations of intercrossing seem to increase the proportion of Westonia alleles. One potential cause for segregation distortion on Chromosome 6B could be the introgression from wild emmer wheat (*Triticum turgidum* ssp. *dicoccoides*) present in the Volcani founder which carries the NAM-B1 gene, also known as GPC-B1 [Uauy et al., 2006, Distelfeld et al., 2012].

**Table 4:** Genetic composition at position 308 cM on chromosome 6B, which has the highest rate of Baxter alleles on chromosome 6B.

| Subpopulation | AC-Barrie | Alsen | Baxter | Pastor | Volcani | Westonia | Xiaoyan | Yitpi |
| --- | --- | --- | --- | --- | --- | --- | --- | --- |
| All | 0.09 | 0.10 | 0.19 | 0.09 | 0.09 | 0.19 | 0.07 | 0.17 |
| MP8RIL | 0.10 | 0.10 | 0.19 | 0.09 | 0.10 | 0.18 | 0.08 | 0.17 |
| AIC2RIL | 0.08 | 0.10 | 0.18 | 0.09 | 0.07 | 0.20 | 0.10 | 0.18 |
| AIC3RIL | 0.08 | 0.11 | 0.19 | 0.07 | 0.09 | 0.23 | 0.05 | 0.18 |

Other localised regions of segregation distortion occur on chromosomes 2D, 4A and 7D. The most distorted point on chromosome 2D occurs at 155 cM; this is likely due to a genetic interaction (discussed in the next seciton). Table S8 gives the genetic composition at 155 cM. The most distorted point on chromosome 4A occurs at 136 cM; Table S9 gives the genetic composition at this point. The most distorted point on chromosome 7D occurs at 72 cM; Table S10 gives the genetic composition at this point. The effect on chromosome 7D may be due to a funnel effect; see the discussion of segregation due to funnel effects. There is another region of distortion at the end of chromosome 7D, at 238 cM. The genetic composition at this second position on chromosome 7D is substantially different to the composition at the first distorted position. There is also evidence for distortion on chromosomes 1D and 5A.

The P-values for a test for distortion are numerically equal to 0 for the localised distortions on chromosomes 2D, 4A and 7D. For localised distortion, P-values should be interpreted extremely cautiously. They may, for example, indicate mapping or genotyping errors, which invalidate the underlying assumptions [Greenland et al., 2016]. Our use a genotyping error rate parameter in haplotype probability computation and imputation tends to guard against these types of errors.

#### Pairwise effects

Interactions were identified as “significant” if the P-value was smaller than 10^−9.5^; this fairly strict threshold was chosen to account for multiple hypothesis testing. There are two significant interactions.

The first is between position 445 cM on chromosome 2B and position 127 cM on chromosome 2D. Although we give the most significant position, the range of locations for which this interaction is significant is very large, especially on chromosome 2B. A table showing the most significant 2B-2D interaction is given in Table S12. A visual representation of the interaction is shown in Figure S13, where colour represents the observed frequency as a multiple of the expected frequency under independence. Dark blue represents combinations present much more frequently than expected under independence. We see that lines with Baxter alleles at both locations occur much more frequently than expected. We also see that lines with a Baxter allele on chromosome 2B and a Xiaoyan allele on chromosome 2D occur much less frequently than expected. We note that the segregation distortion detected at 155 cM on chromosome 2D may be caused by the interaction term detected here.

This interaction might be due to an issue with marker calling for markers on chromosomes 2B and 2D. The large segment of Timopheevi intrgrogression (most of the long arm of 2B) means that the Aestivum 2B chromosome is missing. So the normal hybridisation state for many 2D markers will appear to have a lower copy number and potentially a theta shift. The interaction might also be due to an issue with the map construction process, with some markers being mapped to the wrong chromosome. However, an extensive search for mapping errors failed to identify problems that could be responsible for this interaction.

The second significant interaction is between chromosomes 2B and 6B, and occurs over a more limited region. The location on chromosome 6B (306 cM) is almost the same as one of the locations identified for a main effect, so a genetic interaction may be the cause of that main effect. Table S13 shows the joint distribution of the underlying alleles at position 306 cM on chromosome 6B and position 462 cM on chromosome 2B. A visual representation of the interaction is shown in Figure S14.

The nominal significance threshold of 10^−9.5^ is extremely conservative, and there are likely to be other interaction terms. This conversatism is necessary because, in our experience, markers polymorphic on multiple chromosomes can introduce an erroneous (yet highly significant) interaction. Using the Bonferroni correction to account for the 22,992,937 tests results in an adjusted significance threshold of 0.0073. The Bonferroni correction is likely to be extremely conser-vative in this case. Use of P-values assumes correctness of the genetic map and the marker calling process; P-values should be used with caution.

#### Sex-specific effects

A plot of the chi-squared test statistics for the presence of a sex-specific effect is shown in Figure S16, for every founder and every position. There is one very clear effect on chromosome 6B, related to the sex of the Westonia line in the initial cross. The most significant position for this effect is 158 cM. At this point, the Westonia allele is present in 19% of lines. For lines where Westonia is always a maternal contribution, the Westonia allele is present in 9% of lines. For lines where Westonia is always a paternal contribution, the Westonia allele is present in 29% of lines. The nominal P-value of this effect (ignoring multiple testing) is 8.32 × 10^−10^.

#### Funnel effects

Next, we consider effects relating to the specific combinations of founders in the original cross (funnels). A plot of the chi-squared test statistics for every founder is shown in Figure S17. There is an obvious effect relating to founder Volcani on chromosome 6B, which appears to occur chromosome-wide. Table 5 shows the effect at position 0 cM, which is one of the places where the P-value of the effect is numerically zero. If founder Volcani is crossed with Westonia, Xiaoyan or Yitpi in the first generation, then the haplotype on 6B is extremely unlikely to contain any Volcani alleles. If Volcani is crossed with Baxter in the initial cross, then inheritance of Volcani alleles on 6B is inflated. Table 5 understates the size of the effect, particularly with respect to Westonia / Volcani crosses. Out of 355 MP8RIL lines that cross Westonia and Volcani in the first generation, only 1 has greater than 0.2 probability of having a Volcani allele at 0 cM. It is possible that no lines from these crosses carry a Volcani allele at this position.

**Table 5:** Proportion of Volcani alleles present at location 0 cM on chromosome 6B, if Volcani is crossed with the specified founders in the first generation. If Volcani is crossed with Westonia, Xiaoyan or Yitpi in the initial cross, then the haplotype at 0 cM on chromosome 6B is highly unlikely to be contributed by Volcani.

| AC-Barrie | Alsen | Baxter | Pastor | Westonia | Xiaoyan | Yitpi |
| --- | --- | --- | --- | --- | --- | --- |
| 0.12 | 0.15 | 0.18 | 0.10 | 0.01 | 0.02 | 0.03 |

As the Volcani funnel effect on chromosome 6B occurs chromosome-wide, it also occurs at the location on chromosome 6B where we previously detected segregation distortion (308 cM), and at the location where we previously detected an interaction term (306 cM). It is possible all three of these effects have the same underlying genetic cause.

There are two slightly less significant effects relating to Pastor. The first is at 299 cM on chromosome 3B, with a nominal P-value of 3.1 × 10^−13^. Table 6 shows the chromosome 3B effect; if Pastor is crossed with Baxter or Yitpi in the first generation, then it is unlikely to find a Pastor allele at 299 cM on chromosome 3B. Out of 341 MP8RIL lines that cross Pastor and Yitpi in the first generation, only 5 lines have greater than 0.5 probability of having a Pastor allele at 299 cM. Out of 332 MP8RIL lines that cross Pastor and Baxter in the first generation, only 2 lines have greater than 0.5 probability of having a Pastor allele at 299 cM.

**Table 6:**
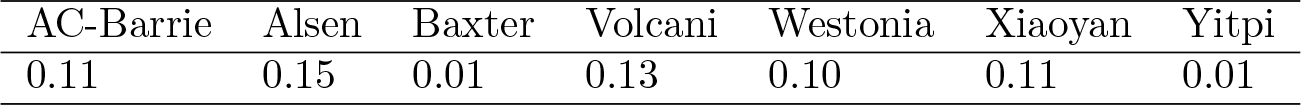
Proportion of Pastor alleles present at location 299 cM on chromosome 3B, if Pastor is crossed with a specific founder in the first generation. If Pastor is crossed with Baxter or Yitpi in the initial cross, then the haplotype at 299 cM on chromosome 3B is highly unlikely to be contributed by Pastor.

| AC-Barrie | Alsen | Baxter | Volcani | Westonia | Xiaoyan | Yitpi |
| --- | --- | --- | --- | --- | --- | --- |
| 0.11 | 0.15 | 0.01 | 0.13 | 0.10 | 0.11 | 0.01 |

There is another effect relating to Pastor on chromosome 7D at position 12 cM, with a nominal P-value of 8.0 × 10^*−12*^. Table 7 shows the chromosome 7D effect; if Pastor is crossed with Baxter or Yitpi in the first generation, then it is unlikely to find a Pastor allele at 12 cM on chromosome 7D. Out of 341 MP8RIL lines that cross Pastor and Yitpi in the first generation, only 4 lines have greater than 0.5 probability of having a Pastor allele at 12 cM. Out of 332 MP8RIL lines that cross Pastor and Baxter in the first generation, only 3 lines have greater than 0.5 probability of having a Pastor allele at 12 cM.

**Table 7:** Proportion of Pastor alleles present at location 12 cM on chromosome 7D, if Pastor is crossed with a specific founder in the first generation. If Pastor is crossed with Baxter or Yitpi in the initial cross, then the haplotype at 12 cM on chromosome 7D is highly unlikely to be contributed by Pastor.

| AC-Barrie | Alsen | Baxter | Volcani | Westonia | Xiaoyan | Yitpi |
| --- | --- | --- | --- | --- | --- | --- |
| 0.13 | 0.13 | 0.01 | 0.09 | 0.12 | 0.11 | 0.01 |

It is highly unlikely for a funnel effect to arise by chance; most types of errors (e.g. mapping errors, genotyping errors) would not cause such an effect. So the effects on chromosomes 3B and 7D represent *at least one* funnel effect. But the funnel effects on chromosomes 3B and 7D appear very similar; it is concievable that there is only a *single* effect which, due to a mapping error, (wrongly) appears in two different regions. The most significant interaction between positions 295 cM - 303 cM on chromosome 3B and positions 8 cM - 16 cM on chromosome 7D has a P-value of 0.24. This suggests that there may be two separate effects.

### Heterozygosity

Figure S18 shows the distribution of the proportion of residual heterozygosity, as determined by the Viterbi algorithm. There are 75 lines with a proportion or residual heterozygosity greater than 0.2. This is likely due to mistakes during the selfing stages of the pedigree, where a line was outcrossed instead of being self-pollinated; it is difficult to guarantee that selfing occurs.

The proportion of residual heterozygosity identified by the Viterbi algorithm was 0.0146 for the MP8RIL lines, 0.034 for the AIC2RIL lines and 0.0273 for the AIC3RIL lines. The theoretical expected residual heterozygosity proportions are 0.0312 without intercrossing, and 0.0273 with intercrossing. While the values for the AIC2RIL and AIC3RIL populations roughly agree with the theoretically expected proportion, the MP8RIL subpulation contains substantially less residual heterozygosity than expected.

## Conclusion

We have presented a large, densely genotyped, high-resolution mapping population and demonstrated its use for a variety of investigations providing new insight into genomic structure in wheat. We have validated a high-density map with the highest resolution currently available in any single wheat population, and used it to identify recombination breakpoints and characterize regions of segregation distortion.

We have identified numerous regions which display distortion in a general sense. In some cases this is due to introgressions (e.g., Chr. 2B). Chromosome 6B is particularly interesting, as we find evidence for segregation distortion, an interaction term, a sex-specific effect *and* a funnel effect. It is unclear how many underlying genetic causes there are for these effects; potentially, there could be a single causal locus, however the effects identified could have significant implications on breeding programs selecting for the GPC locus on 6B. Identification of these types of genetic effects is vital for translational research in crops, as it leads to a deeper understanding of genomic structure and its influence on breeding decisions. This population provides a high-value genetic resource useful for better understanding basic wheat genetics and improving the crop.

In the map presented in this paper, we have excluded markers that are polymorphic on more than one chromosome. However, it is feasible to map such markers to multiple locations. As the positions of these markers on different chromosomes are likely to be related, they are particularly useful for understanding homeoallele effects and contributions to phenotypes. It is also possible to use the constructed genetic map to identify and call additional marker alleles, for currently mapped markers.

Here we have focused on analysis of genomic structure facilitated by the large number of lines genotyped at high density. This population has undergone phenotyping for a large number of traits, which almost always display transgressive segregation. In future studies we hope to gain some insight into the effect of segregation distorting genetic effects on phenotypic performance.

By building on the foundation presented here with further investigation of how the genomic structure contributes to phenotypic diversity, we expect that our understanding of the importance of structural diversity in wheat can be better understood and utilised for wheat improvement.

## Methods

### Population

The population is divided into the three subpopulations shown in Figure 3. For each subpopulation the pedigree has three stages, as described in [Valdar et al., 2006]. In the first stage, mixing, the parent genomes are combined together over three generations of mating. In the first generation, 28 crosses were made in a single direction (no reciprocal crosses). In the second generation, 210 crosses were made in a single direction. In the third generation, 589 distinct crosses were created, including reciprocal crosses in 276 cases. All unique combinations of founders, referred to as *funnels* are shown in Figure 2, divided by each stage of the mixing process. The G3 individuals were common to all subpopulations and encompassed 313 of the 315 unique combinations possible for an eight-way population, excluding reciprocal crosses. Figure S19 shows the structure of the funnels, and the level of representation of each funnel at each stage in the final population. In the second stage, maintenance or intercrossing, individuals from different funnels were intercrossed to generate additional recombination between genome segments. There were 293, 286 and 745 lines in the first, second and third generations of intercrossing, respectively. Finally, in the third stage, inbreeding, individuals were selfed for five generations. In all, there were 2151 distinct crosses made to generate the complete population.

**Figure 2:**
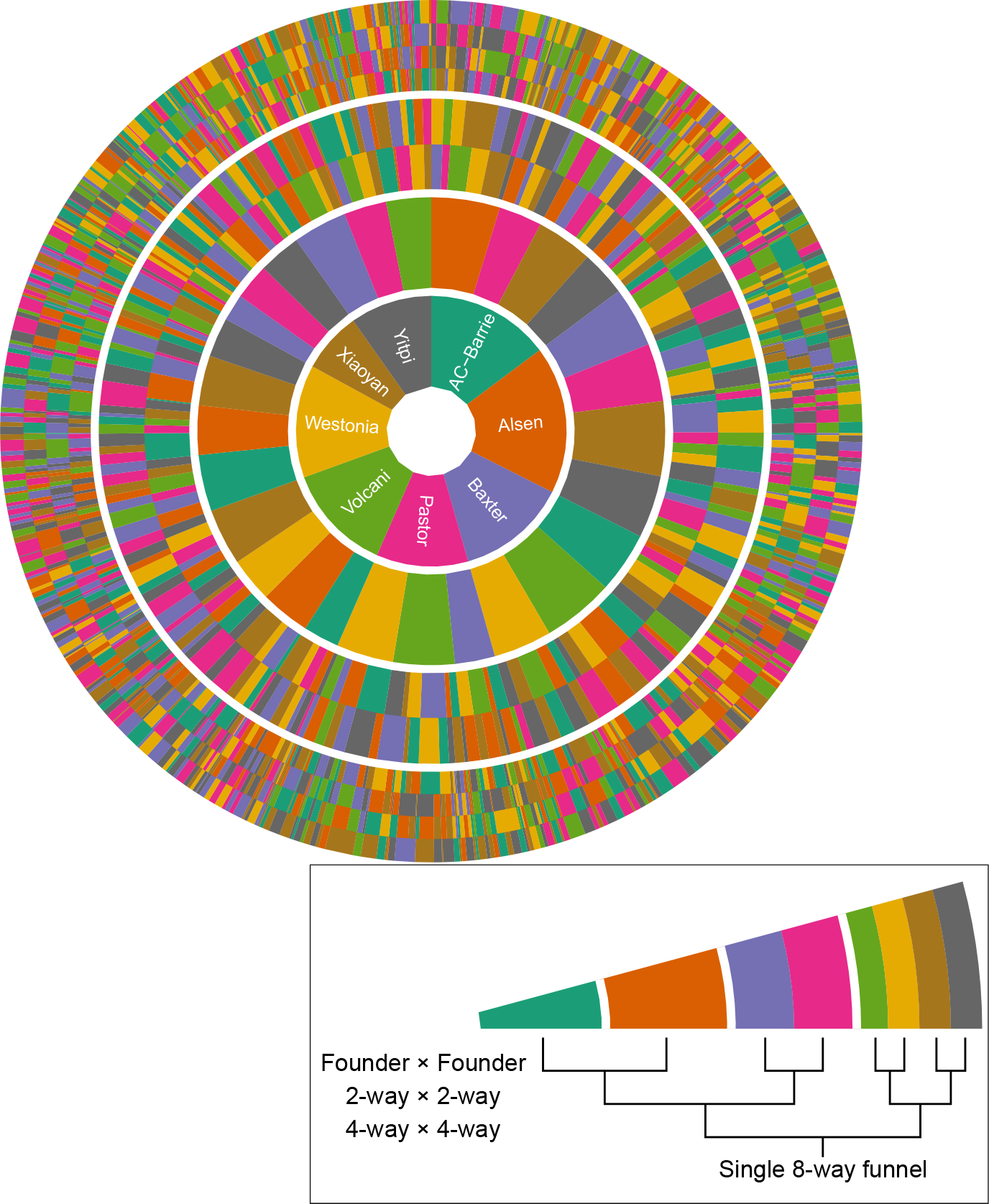
Radial representation of all 559 funnels used to generate the M8 RILs. An example single funnel is shown in the inset. Each ring in the figure represents a plant in each stage of the crossing strategy. The first, most central ring is the maternal founder in the first round of crossing. The second ring is the paternal founder in the first round. The third ring shows the founder makeup of the paternal 2-way line used in the second round of crossing. The outermost ring shows the founder makeup of the paternal 4-way line in the third round of crossing.

**Figure 3:**
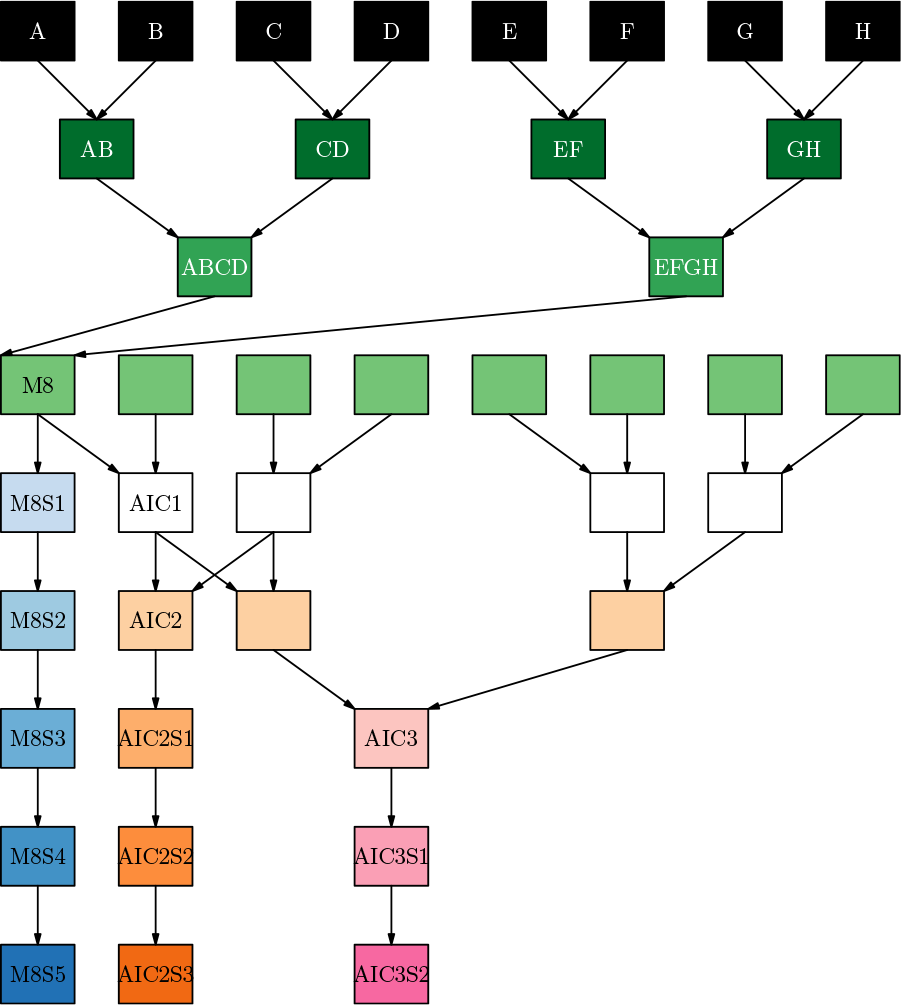
Schematic of the different branches of the three sub-populations. Genotyped individuals are shown in black. Levels indicate generations of crossing; individuals of the same color have undergone similar breeding up to that stage of the pedigree; S* indicates generation of selfing. M8 lines undergo only mixing of the parents and inbreeding; AIC2 and AIC3 lines undergo two and three additional generations of intercrossing prior to inbreeding, respectively. This schematic represents a single funnel for each sub-population type.

The first subpopulation (MP8RIL) contained 2381 lines generated using mixing and selfing, but no generations of intercrossing. Of the possible 315 unique funnel combinations, 311 were used in the MP8RIL subpopulation. The second subpopulation (AIC2RIL) contained 286 lines generated using two generations of intercrossing prior to inbreeding. The third subpopulation (AIC3RIL) contained 745 lines generated using three generations of intercrossing prior to inbreeding.

### Genotyping and map construction

A preliminary map was constructed using five steps; genotype calling, recombination fraction estimation, linkage group identification, marker ordering, and map estimation. These steps were performed using R packages **mpMap2**, **mpMapInteractive2**and **magicCalling**. Package **mpMap2**[Shah and Huang, 2019] is an R package for map construction using multi-parent crosses, and is an updated version of **mpMap**[Huang and George, 2011]. Package **mpMapInteractive2**[Shah, 2019] is a package for interactively making manual changes to linkage groups and marker ordering, during the map construction process. Package **magicCalling**contains code used for SNP calling. Unless otherwise noted, all R functions are contained in **mpMap2**.

After the preliminary map was constructed, the map was improved incrementally, until a final map was constructed, and it is this final version that is discussed in this paper. It is not possible to construct a genetic map in a single pass, as some types of improvements can only be made *after* a preliminary map has been constructed. For example, some marker calling errors cannot be accurately identified without a genetic map; see Figures S3a-S3c for an example.

### Genotype calling

Each marker was first processed using the data normalization approach of GenCall [Peiffer et al., 2006], and then converted to the polar coordinates (*r, θ*). Markers were called using either DBSCAN [Ester et al., 1996] or the hierarchical Bayesian clustering (HBC) model described in [Shah and Whan, 2018]. The HBC model is implemented using the JAGS library [Plummer, 2015]. It has the advantage of calling heterozygotes, but the disadvantage that it cannot be applied if there are more than two marker alleles. This can occur if there are secondary polymorphisms at the target location, or if a marker is polymorphic at multiple locations on the genome. We chose parameters for the HBC method that resulted in fairly aggressive calling of heterozygotes.

DBSCAN can identify more than two marker alleles for a single marker. However it cannot identify heterozygotes, and has two parameters (minPts and *ε*) that must be specified manually for each marker.

These methods have advantages over the GenCall algorithm implemented in Illumina GenomeStudio; DBSCAN can call more than two marker alleles, while the HBC approach can call heterozygotes.

We performed genotype calling by first applying HBC to every marker. We identified monomorphic markers, and lines that had a high error rate. These lines were removed, and HBC was reapplied to the polymorphic markers. A subset of the fitted HBC models were reviewed manually, based on some simple heuristics. Reasons for needing to review a marker included non-convergence of the MCMC algorithm used to fit the HBC model, an unreasonably high rate (>0.06) of heterozygote calls, or the presence of more than two marker alleles. If more than two marker alleles were found, DBSCAN was used to define clusters. Both DBSCAN and the HBC model are implemented in the **magicCalling** package.

### Estimation of recombination fractions

Recombination fractions between all 437,059,395 pairs of called markers were estimated using the function estimateRF. Estimation was performed using numerical maximum likelihood, with 61 possible recombination fraction values. This step took 14 hours, and required 300GB of memory. Chromosome 2B carries the Sr36 introgression from Triticum timopheevi [Tsilo et al., 2008], which is known to distort genetic inheritance. This was corrected for in later steps, using the weighting method described in [Shah et al., 2014]. The weight assigned to a particular line depended not only on whether the introgression was present or absent, but the number of generations of intercrossing.

### Construction of linkage groups

The set of all markers was divided into 400 smaller groups, by applying hierarchical clustering to the matrix of recombination fractions, using the function formGroups. The underlying implementation comes from the **fastcluster** package [Müllner, 2013]. These small groups were then aggregated by hand using **mpMapInteractive2**, resulting in 29 groups of appreciable size (> 10 markers). The linkage groups corresponding to the 21 chromosomes were identified based on the consensus map [Wang et al., 2014]. In some cases it was difficult to identify the correct linkage group for a chromosome, especially for the D chrom-somes.

### Marker ordering

Ordering of chromosomes proceeded in three steps. The first step was performed using the clusterOrderCross function. Each chromosome was divided into 30 subgroups using hierarchical clustering. These subgroups were used to define a 30 × 30 matrix, where each entry was the average of the recombination fractions between markers in those subgroups. The 30 subgroups were automatically ordered by applying anti-robinson serialization [Brusco et al., 2007, Hahsler et al., 2008] to this 30 × 30 matrix. At the end of this step, the ordering of markers *within* a subgroup is still arbitrary. An example of this step is shown in Figures S20a-S20c.

The second step was the ordering of markers within chromosomes, using the orderCross function. This function applies a modified version of anti-robinson serialization to the matrix of recombination fractions. Standard anti-robinson serialization allows global changes to the marker ordering. The modified version only makes local changes to the ordering, and is therefore computationally much faster. The result of this step is shown in Figure S20d.

The third step was manual changes made using the **mpMapInteractive2** package, based on visual inspection of the recombination fraction heatmaps.

### Map distance estimation

Map distance estimation was performed by forming a collection of equations, and approximately solving them using non-linear least squares. These equations allow the genetic distances between pairs of markers, which are not adjacent in the chosen ordering, to be used as part of the estimation process. Consider the case where there are three markers, known to be in the correct order. If the matrix of estimated genetic distances (obtained from the estimated recombination fractions) is

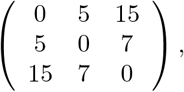

then the matrix equation to be solved is

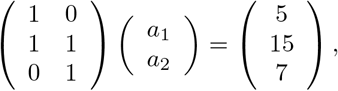

where *a*_1_ is the distance between the first and second markers, and *a*_2_ is the distance between the second and third markers. This method of estimating the genetic map is implemented by the `estimateMap` function. See [Shah and Huang, 2019] for further details.

### Incremental changes

After an initial map was constructed, it was improved incrementally, resulting in the map presented in this paper. Incremental changes are necessary, as some improvements are only possible *after* a preliminary map has been produced. Three types of incremental changes were made. The first was the deletion of markers that were found to be badly called (based on visual inspection of the HBC models), or polymorphic on multiple chromosomes. The second was the addition of markers that could be localised to specific genetic regions, using the preliminary genetic map and a qtl-mapping approach, with marker data being used as a trait.

The third was the identification of ‘dangling’ linkage groups, as being part of a specific chromosome. In particularly marginal cases, we relied on the the International Wheat Genome Sequencing Consortium (IWGSC) RefSeq v1.0 sequence.

### Recombination

Recombination events were imputed by assuming a hidden Markov model (HMM) for the identity-by-descent genotypes. This assumption is not exact, but is known to be highly accurate as long as the distances between consecutive markers are not large. The HMM assumption allows the application of the Viterbi algorithm to impute the most likely underlying genotype, for each line, including imputation of heterozygosity. The Viterbi algorithm is implemented in **mpMap2**. We used an error parameter of 0.1. The underlying probability computations build on previous published work [Teuscher and Broman, 2007, Broman, 2012].

### Segregation Distortion

We ignored marker heterozygotes throughout our asessment of segregation distortion.

#### Main effects

We performed tests for segregation distortion at a locus as follows. For each potential genotype at that position, we computed the average, over all genetic lines, of the probability that this genotypes occured in each line. These values are then arranged into a vector, and the standard chi-squared test for independence was performed.

This statistic is generated from a “contingency table” of non-integer values, however it is still a valid test (central limit theorem). We applied this test for all genetic locations, using a 1 cM grid of equally spaced points.

#### Pairwise effects

Genomic incompatibilities between parents can be detected by further analysis of segregation distortion on the level of interactions. We performed tests for the presence of interactions between a pair of genetic locations as follows. For each potential combined genotype at these positions, we computed the average, over all genetic lines, of the probability that these genotypes occured in each line. These values are then arranged into a 8 × 8 matrix, and the standard chi-squared test for independence was performed. An example of the matrix used in this test is shown in Table S12. The top-left value is the average over all genetic lines of *P* (*L*_1_ = AC-Barrie*, L*_2_ = AC-Barrie), where *L*_1_ and *L*_2_ are genetic locations. In this case the value is around 0.01.

This statistic is generated from a “contingency table” of non-integer values, however it is still a valid test. We applied this test for all pairs of genetic locations on different chromosomes, using a 1 cM grid of equally spaced points.

#### Sex-specific effects

We also investigated segregation distortion that occurs in specific cross combinations (funnels). For each line in the MP8RIL subpopulation, one founder was maintained as a maternal contribution in every cross, and one was maintained as a paternal contribution in every cross. For each founder *f*, we took two subsets of the MP8RIL population - those which had *f* as only a maternal contribution, and those which had *f* as only a paternal contribution. For each genetic location on the 1 cM grid considered previously, we performed a chi-squared test for the presence of a sex-specific effect on the inheritance of *f*. Similar to the interactions, these tests were based on sums of haplotype probabilities.

#### Funnel effects

Another cause of segregation distortion is potential incompatibilities between pairs of founders in the initial cross, which may lead to segregation distortion in the final generation. For each founder *f*, we separated the MP8RIL lines into seven groups, based on the founder which was crossed with *f* in the first generation. We ignored the sex of *f* when forming these groups. We then performed a chi-squared test for differences in genetic inheritance of *f* between these seven groups. This tests the presence of a funnel-specific effect, for the inheritance of founder *f*. Similar to the interactions, these tests were based on sums of haplotype probabilities.

## Declarations

### Availability of data and materials

The genetic map, along with the genetic data for the population, and imputed genotypes, are available online [Shah et al.]. This analysis was performed using packages **mpMap2** [Shah and Huang, 2019], **mpMapInteractive2** and **magicCalling** [Shah and Whan, 2018], which are all publically available [Shah, 2018b, 2019, 2018a] from Github.

### Authors contributions

CC and MKM conceived the study. CC and MN developed the population. RS and BEH constructed the genetic map and performed analysis of segregation distortion. KV performed QTL analysis. RS, BEH and AW wrote the manuscript, and all authors read and approved the final manuscript.

## Acknowledgements

The authors wish to acknowledge the hard work and dedication from a team of people who made such a complex project possible with their tireless effort and attention to detail. Whilst there were many contributors we would like to pay special thanks to Zena Nath, Ibrahim Ibrahim Kutty, Lynnette Rampling, Zoe Campbell, Aswin Singaram Natarajan, Alexandre Boyer, Mark Cmiel, Patrick Moody, Jackson Tahapehi, AJE Haddis, Tracy Willis, Emmett Leyne, May Sun and Kathy Dibley.

## Ethics approval and consent to participate

Not applicable.

