## Supplementary Figures and Tables for "The complex genetic architecture of recombination and structural variation in wheat uncovered using a large 8-founder MAGIC population"

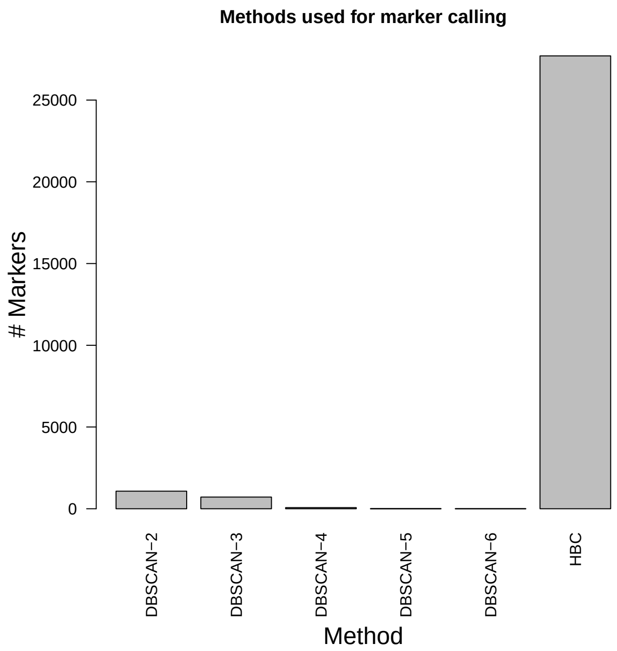


Supplementary Figure 1: Number of times each genotype calling method was used. Method “DBSCAN-n” indicates the use of DBSCAN, where $n$ is the number of alleles identified by DBSCAN.

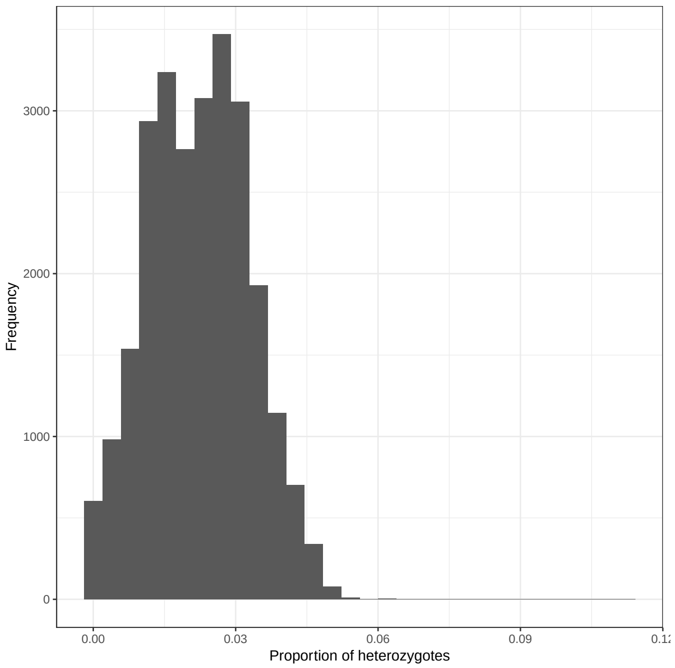


Supplementary Figure 2: Distribution of the proportion of *marker* heterozygotes, for markers called using HBC. Note that not all identity-by-descent heterozygotes are marker heterozygotes.

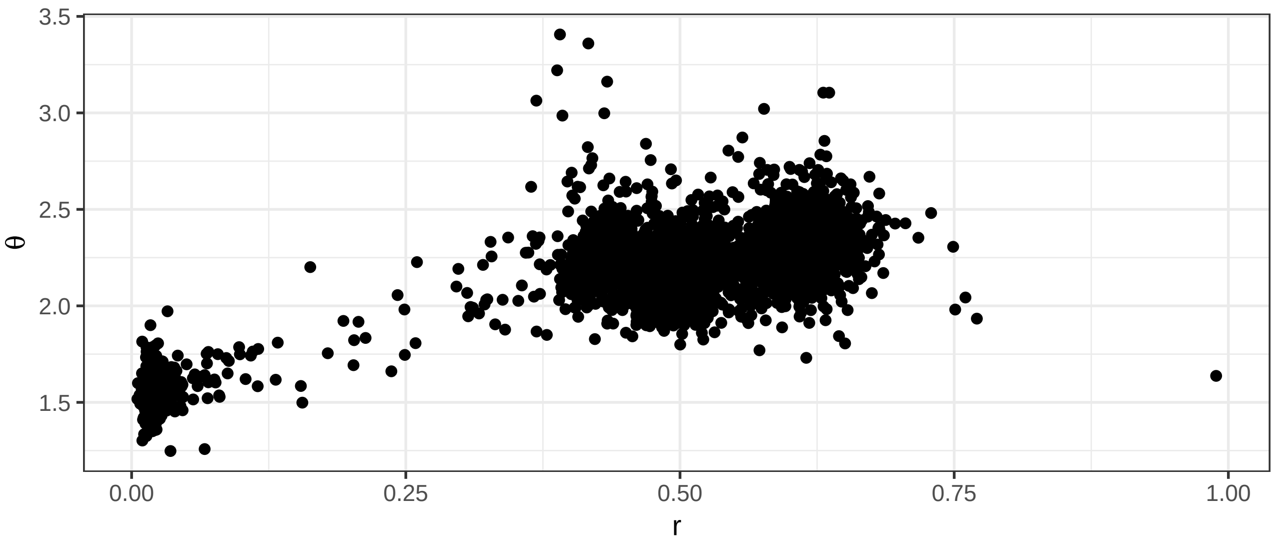


(a) Marker data, without a genetic map.

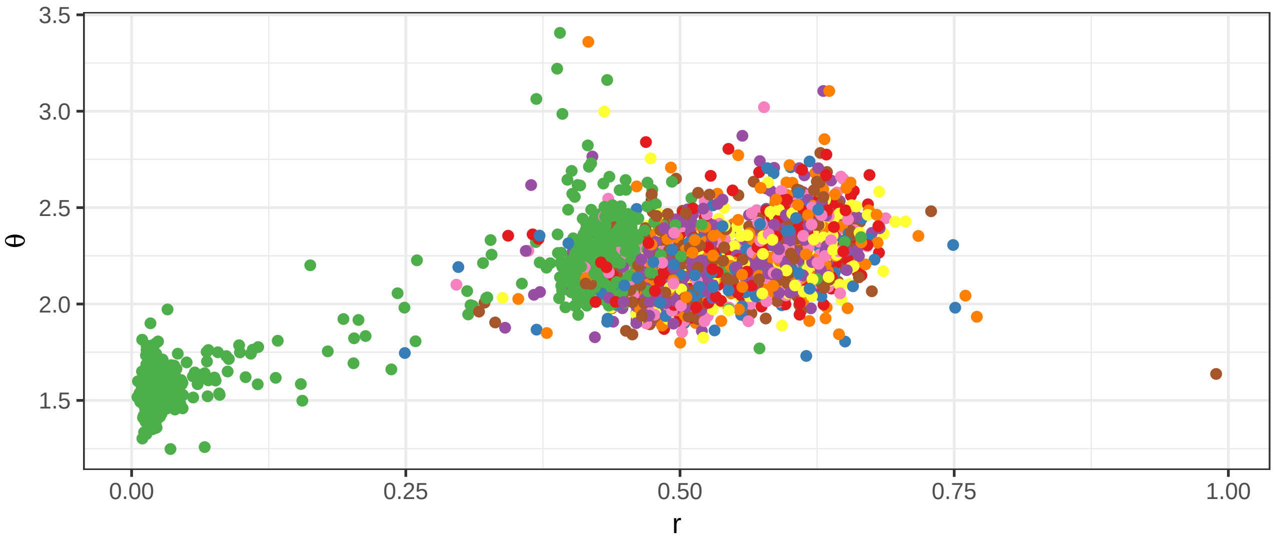


(b) Plot of the marker data, after a genetic map is constructed. Color represents the final imputed founder genotype, at a location on chromosome 2B.

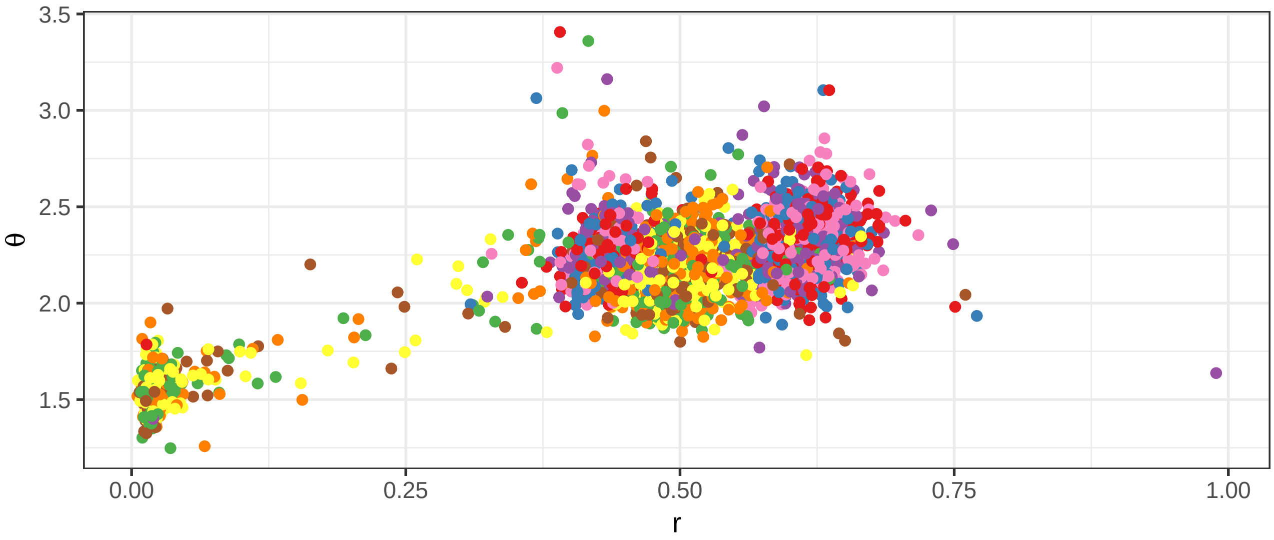


(c) Plot of the marker data, after a genetic map is constructed. Color represents the final imputed founder genotype, at a location on chromosome 2A.

Supplementary Figure 3: Example of a marker that is polymorphic on more than one chromosome. This marker (Kukri_c37840_253) is polymorphic on chromosomes 2A and 2B, although this is not obvious without a genetic map.

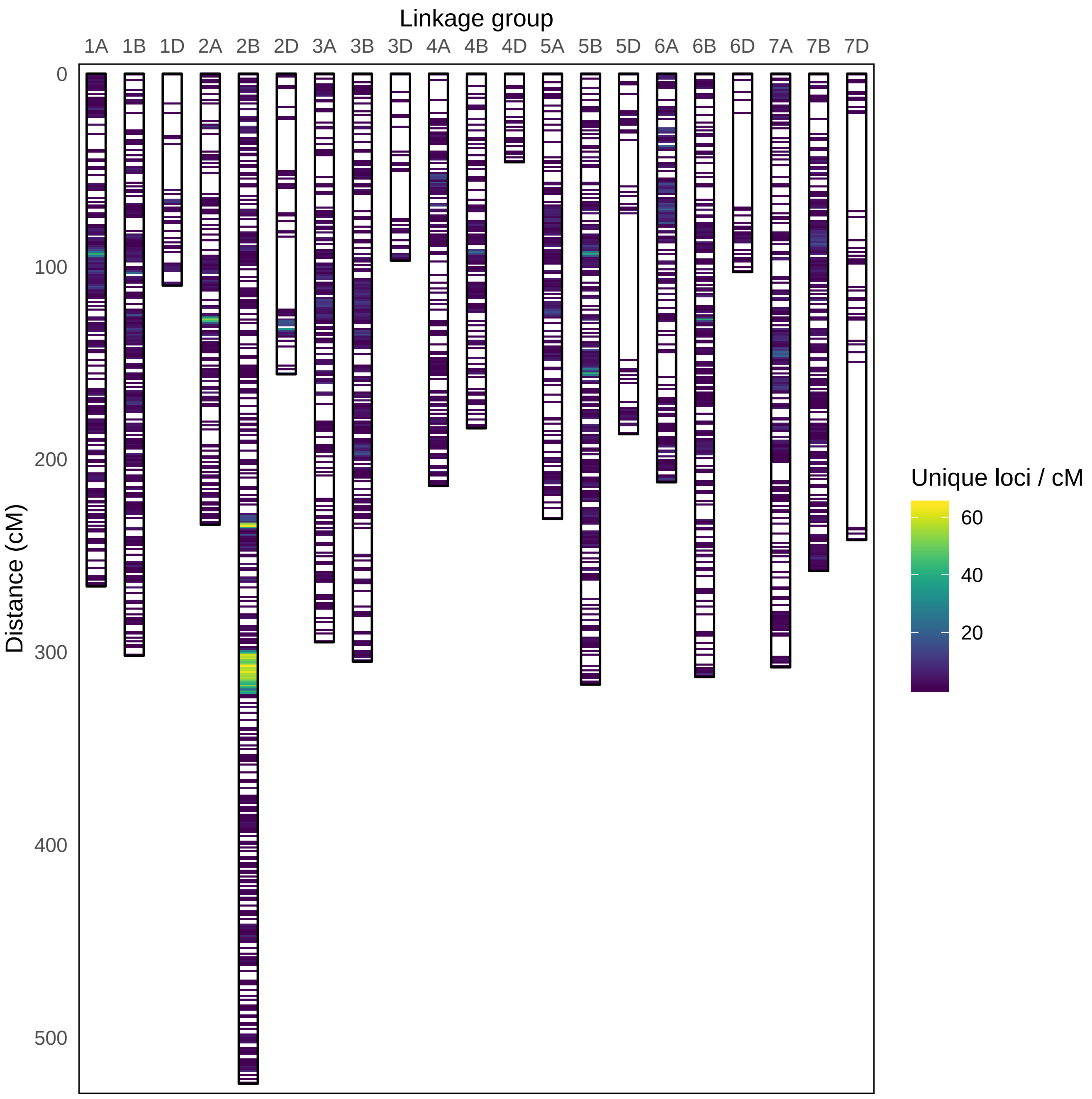


Supplementary Figure 4: Illustration of the final map generated from the MAGIC population. Color represents unique loci / cM.

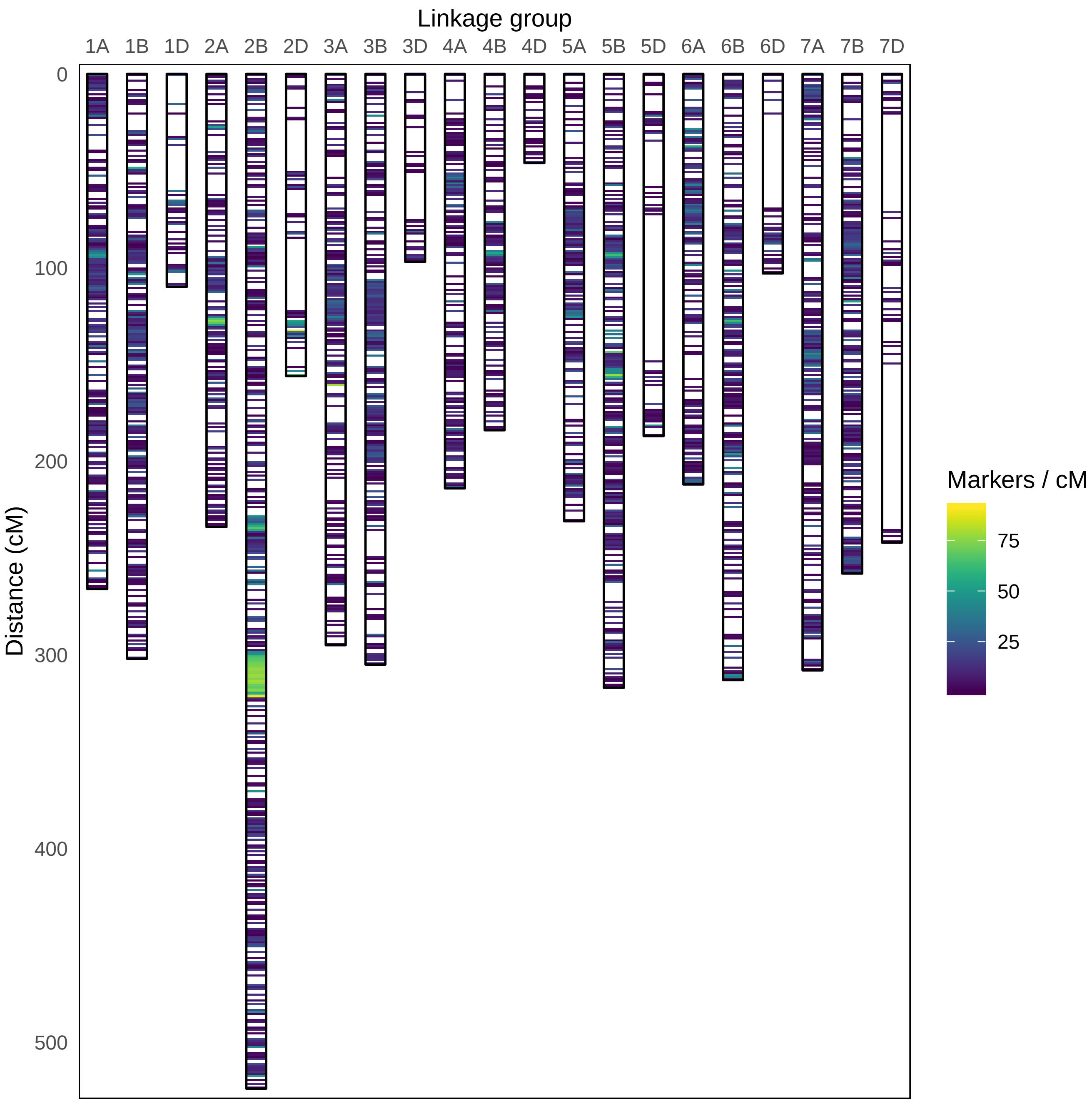


Supplementary Figure 5: Illustration of the final map generated from the MAGIC population. Color represents markers / cM

| Chromosome | # Markers | # Distorted Markers | # Unique Positions | Length | Resolution |
|---|---|---|---|---|---|
| 1A | 1717 | 114 | 503 | 265.49 | 1.89 |
| 1B | 1988 | 157 | 482 | 301.72 | 1.60 |
| 1D | 393 | 66 | 84 | 109.85 | 0.76 |
| 2A | 1376 | 520 | 389 | 233.54 | 1.67 |
| 2B | 4899 | 3839 | 1886 | 523.63 | 3.60 |
| 2D | 646 | 105 | 171 | 155.96 | 1.10 |
| 3A | 1387 | 73 | 347 | 294.39 | 1.18 |
| 3B | 1832 | 130 | 463 | 304.99 | 1.52 |
| 3D | 148 | 5 | 46 | 96.88 | 0.47 |
| 4A | 1059 | 83 | 269 | 213.08 | 1.26 |
| 4B | 782 | 77 | 208 | 183.68 | 1.13 |
| 4D | 79 | 15 | 25 | 45.93 | 0.54 |
| 5A | 1311 | 238 | 321 | 230.25 | 1.39 |
| 5B | 2392 | 210 | 634 | 316.14 | 2.01 |
| 5D | 244 | 17 | 64 | 186.17 | 0.34 |
| 6A | 1545 | 59 | 453 | 211.40 | 2.14 |
| 6B | 1805 | 998 | 350 | 312.70 | 1.12 |
| 6D | 222 | 22 | 36 | 102.27 | 0.35 |
| 7A | 1854 | 408 | 485 | 307.20 | 1.58 |
| 7B | 1804 | 107 | 416 | 257.62 | 1.61 |
| 7D | 204 | 44 | 42 | 241.16 | 0.17 |
| All | 27687 | 7287 | 7674 | 4894.06 | 1.57 |

Supplementary Table 1: Summary of the genetic map by chromosome, including number of markers mapped; number of markers with segregation distortion ($p < 10^{-5}$); length of chromosomes (centiMorgans); and resolution of the map, defined as the number of unique positions per centiMorgan.

| | | Consensus | | NIAB | |
|---|---|---|---|---|---|
| Mapped | Chromosome | Common | Conflicts | Common | Conflicts |
| 1717 | 1A | 1464 | 40 | 604 | 22 |
| 1988 | 1B | 1741 | 45 | 1180 | 111 |
| 393 | 1D | 347 | 11 | 251 | 0 |
| 1376 | 2A | 1168 | 48 | 540 | 15 |
| 4899 | 2B | 2314 | 673 | 1066 | 402 |
| 646 | 2D | 564 | 11 | 397 | 21 |
| 1387 | 3A | 1161 | 2 | 723 | 4 |
| 1832 | 3B | 1561 | 34 | 968 | 4 |
| 148 | 3D | 109 | 3 | 71 | 0 |
| 1059 | 4A | 938 | 7 | 347 | 34 |
| 782 | 4B | 665 | 2 | 408 | 5 |
| 79 | 4D | 63 | 1 | 47 | 0 |
| 1311 | 5A | 1067 | 90 | 625 | 9 |
| 2392 | 5B | 2087 | 67 | 1269 | 21 |
| 244 | 5D | 165 | 13 | 154 | 1 |
| 1545 | 6A | 1331 | 5 | 904 | 19 |
| 1805 | 6B | 1542 | 203 | 889 | 133 |
| 222 | 6D | 150 | 1 | 115 | 0 |
| 1854 | 7A | 1490 | 101 | 1126 | 2 |
| 1804 | 7B | 1581 | 31 | 609 | 25 |
| 204 | 7D | 164 | 6 | 58 | 5 |

Supplementary Table 2: Comparison of the MAGIC genetic map with the 90K consensus map and the NIAB map. For each chromosome, we present the number of markers mapped in our map (Mapped); the number of those in common with each comparison map (Common); and the number of disagreements (Conflicts) based on markers with distance >20 cM from the best fit curve to the comparison of map positions. Conflicts may be less accurate for D chromosomes as the smaller number of markers can lead to overfitting of the curve.

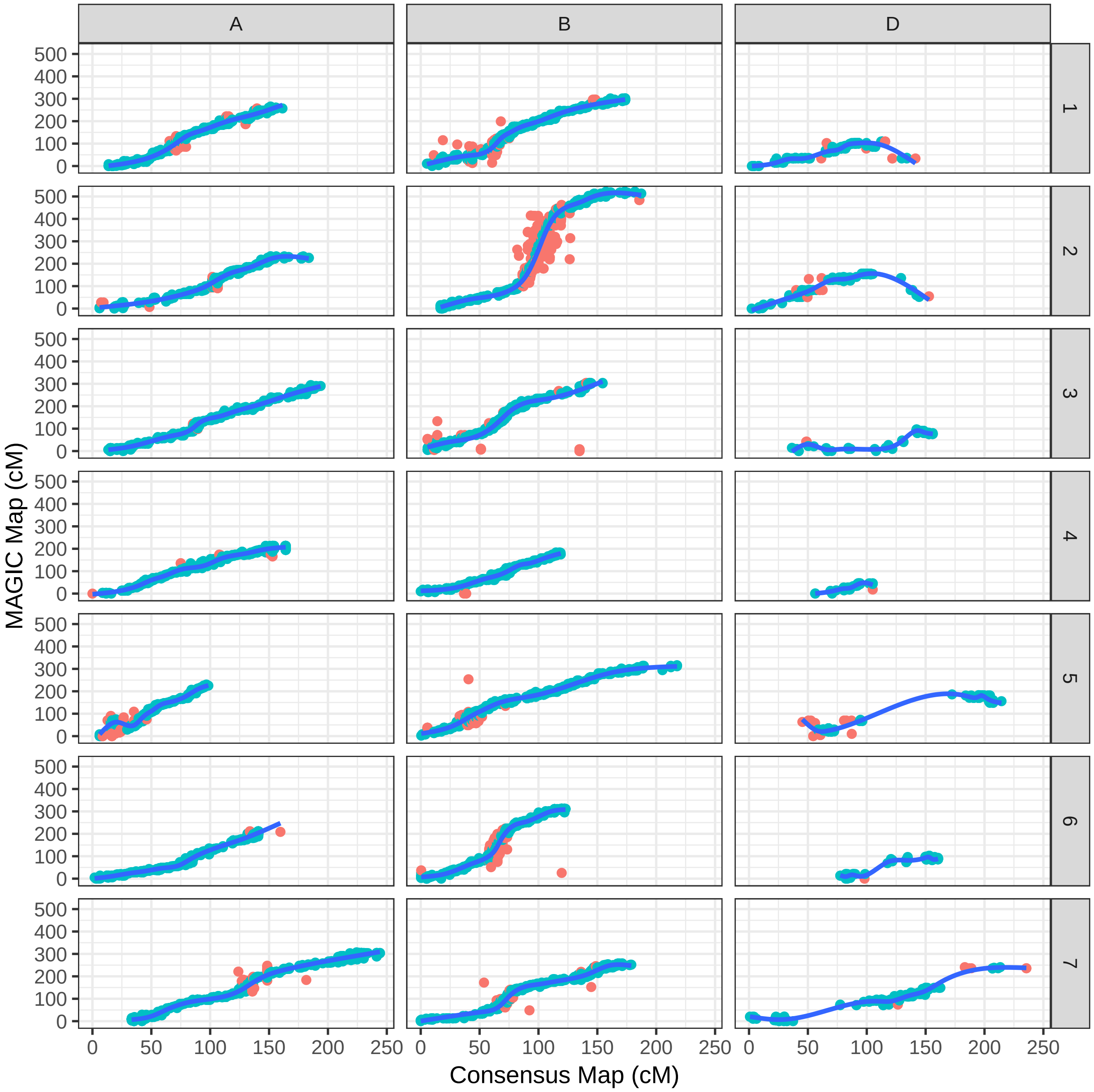


Supplementary Figure 6: Comparison of the MAGIC map with the 90K consensus map. The blue line represents a best fit curve. Red points denote markers whoose positions deviate more than 20cM vertically from the curve.

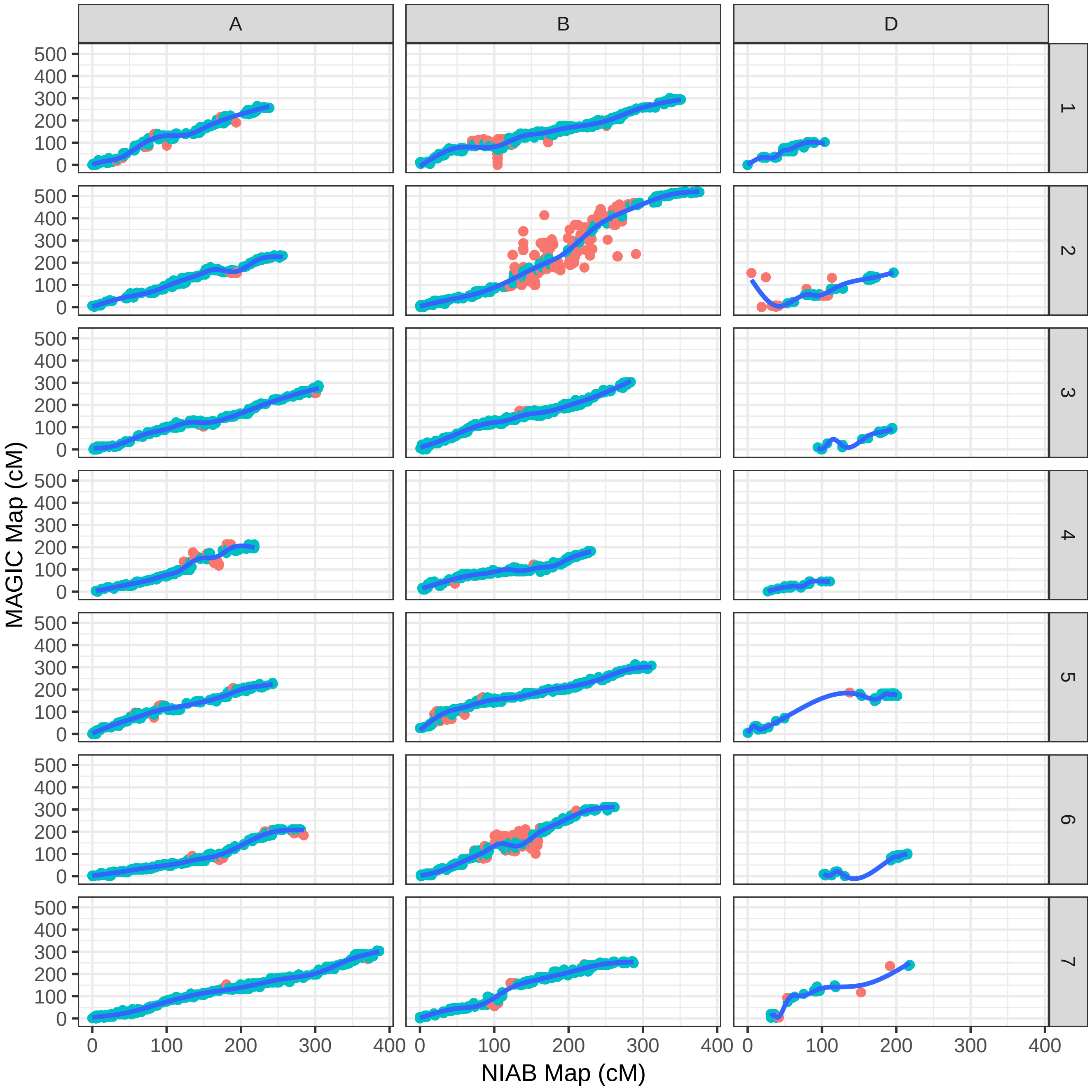


Supplementary Figure 7: Comparison of the MAGIC map with the NIAB map. The blue line represents a best fit curve. Red points denote markers whoose positions deviate more than 20cM vertically from the curve.

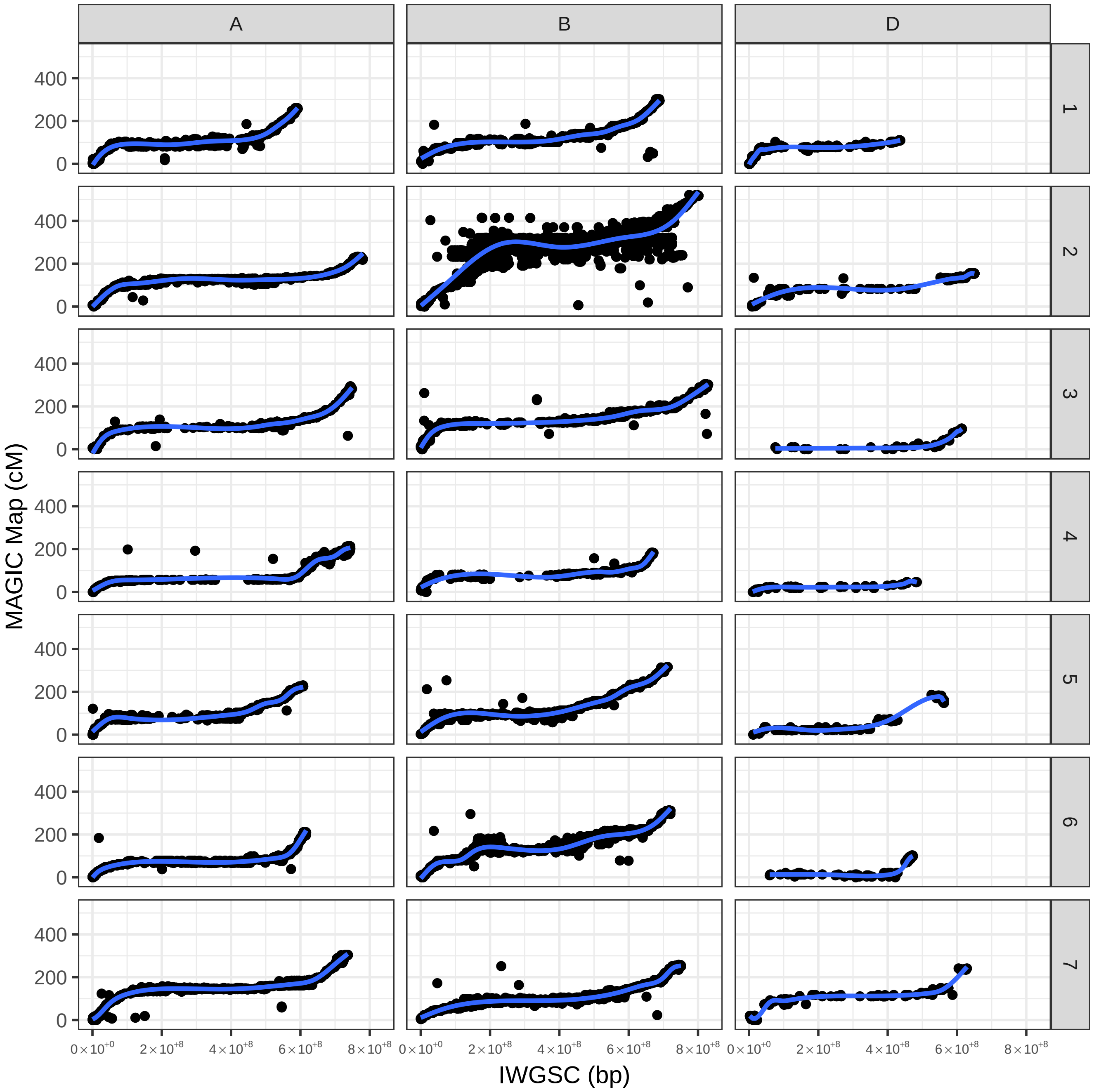


Supplementary Figure 8: Comparison of the MAGIC map with the IWGSC RefSeq v1.0. The blue line represents a best fit curve.

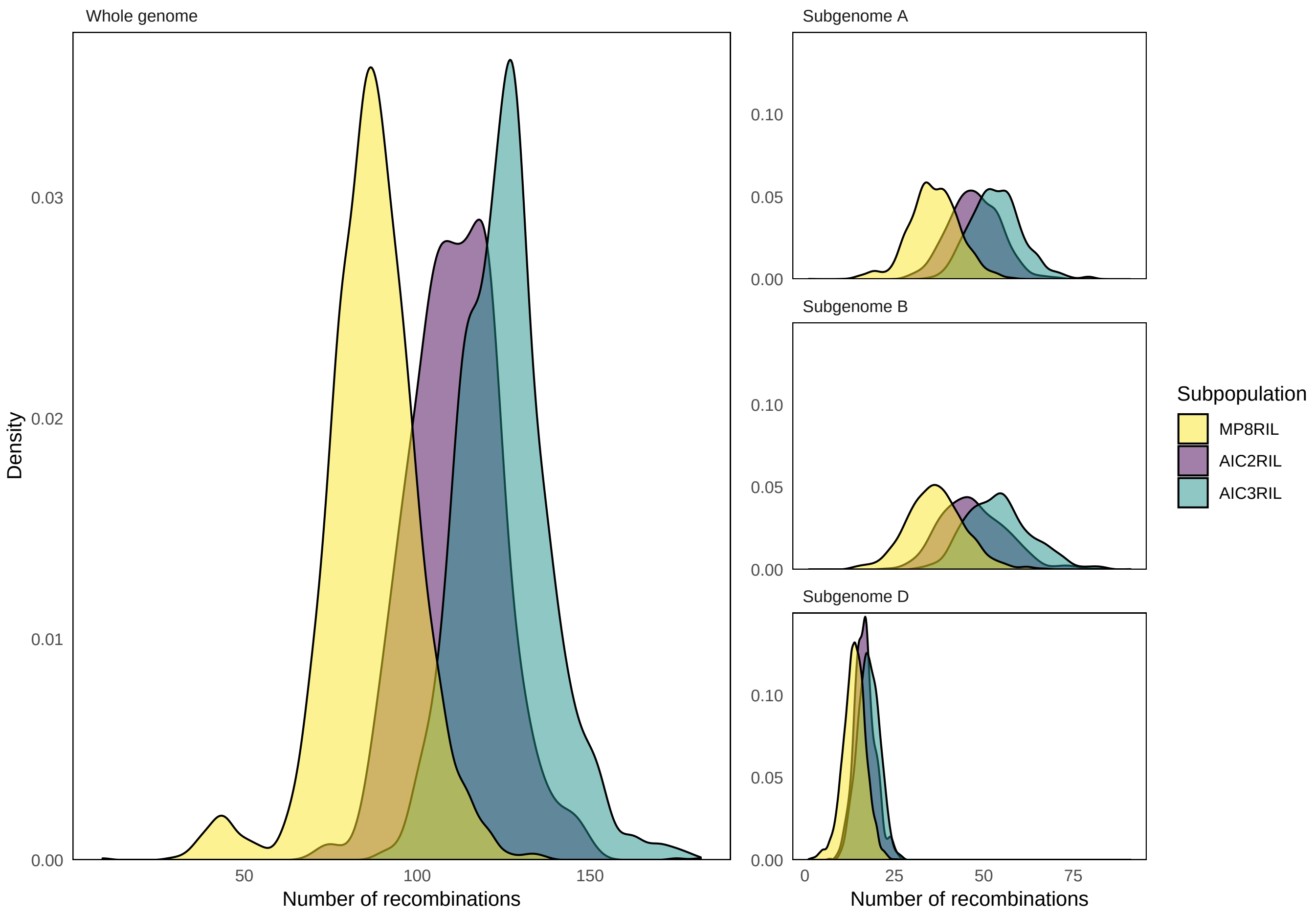


Supplementary Figure 9: Distributions of the total number of recombination events for each subpopulation by, whole genome and subgenomes.

| Chromosome | MPRIL8 | AIC2RIL | AIC3RIL |
|---|---|---|---|
| 1A | 5.15 | 7.09 | 7.80 |
| 1B | 5.26 | 6.75 | 8.10 |
| 1D | 2.18 | 2.79 | 2.81 |
| 2A | 5.01 | 6.71 | 7.37 |
| 2B | 6.62 | 8.30 | 9.40 |
| 2D | 2.64 | 3.05 | 3.37 |
| 3A | 5.85 | 7.64 | 8.90 |
| 3B | 5.67 | 7.42 | 8.40 |
| 3D | 1.57 | 1.92 | 1.91 |
| 4A | 5.03 | 6.61 | 7.46 |
| 4B | 3.52 | 4.80 | 5.21 |
| 4D | 0.77 | 1.02 | 1.16 |
| 5A | 4.79 | 5.79 | 6.60 |
| 5B | 6.93 | 8.87 | 10.39 |
| 5D | 2.24 | 2.74 | 2.95 |
| 6A | 4.19 | 5.35 | 6.15 |
| 6B | 4.16 | 5.34 | 6.22 |
| 6D | 1.42 | 1.69 | 1.80 |
| 7A | 6.20 | 8.21 | 9.23 |
| 7B | 4.59 | 5.76 | 6.56 |
| 7D | 3.01 | 3.51 | 3.77 |
| Whole Genome | 86.82 | 111.36 | 125.57 |

Supplementary Table 3: Average number of recombination events for each chromosome, for each of the three subpopulations.

| Chromosome | Position (cM) | %var | P-value |
|---|---|---|---|
| 1A | 90.71 | 7.10 | 0.03 |
| 1A | 95.63 | 11.89 | 0.00 |
| 1A | 98.73 | 4.95 | 0.04 |
| 1A | 102.53 | 1.66 | 0.34 |
| 1A | 108.41 | 4.11 | 0.00 |
| 1A | 110.54 | 7.82 | 0.00 |
| 2A | 110.42 | 1.26 | 0.00 |
| 3A | 94.31 | 4.04 | 0.00 |
| 3A | 100.42 | 5.26 | 0.00 |
| 4A | 185.22 | 1.14 | 0.00 |
| 4A | 191.24 | 1.05 | 0.00 |

Supplementary Table 4: QTL mapping results, where trait is the number of recombination events on the A subgenome.

| Chromosome | Position (cM) | %var | P-value |
|---|---|---|---|
| 1B | 133.81 | 1.35 | 0.00 |
| 1B | 175.42 | 1.09 | 0.00 |
| 3B | 193.66 | 2.97 | 0.00 |
| 3B | 195.93 | 2.93 | 0.11 |
| 3B | 198.36 | 1.42 | 0.03 |
| 5B | 79.54 | 5.28 | 0.00 |
| 5B | 83.72 | 3.52 | 0.00 |
| 5B | 86.16 | 4.19 | 0.00 |
| 5B | 87.77 | 5.81 | 0.00 |
| 5B | 91.68 | 5.20 | 0.00 |
| 7B | 80.65 | 3.84 | 0.00 |
| 7B | 87.36 | 5.58 | 0.02 |
| 7B | 88.23 | 6.52 | 0.16 |
| 7B | 89.14 | 12.38 | 0.01 |
| 7B | 93.48 | 1.21 | 0.01 |
| 7B | 102.08 | 1.56 | 0.04 |
| 7B | 102.78 | 3.14 | 0.00 |
| 7B | 106.29 | 2.12 | 0.00 |
| 7B | 197.01 | 1.70 | 0.00 |
| 7B | 201.47 | 1.24 | 0.00 |

Supplementary Table 5: QTL mapping results, where trait is the number of recombination events on the B subgenome.

| Chromosome | Position (cM) | %var | P-value |
|---|---|---|---|
| 1D | 48.55 | 1.04 | 0.00 |
| 3D | 50.07 | 1.14 | 0.00 |
| 5D | 150.89 | 1.08 | 0.00 |
| 6D | 78.44 | 9.32 | 0.00 |
| 6D | 83.00 | 5.17 | 0.00 |
| 7D | 9.88 | 1.09 | 0.00 |
| 7D | 149.30 | 1.43 | 0.00 |

Supplementary Table 6: QTL mapping results, where trait is the number of recombination events on the D subgenome.

| Chromosome | Position (cM) | %var | P-value |
|---|---|---|---|
| 2A | 224.40 | 1.97 | 0.00 |
| 2A | 233.29 | 1.13 | 0.02 |
| 2B | 217.02 | 12.38 | 0.00 |
| 2B | 260.76 | 6.69 | 0.00 |
| 2B | 264.14 | 2.56 | 0.02 |
| 5B | 90.49 | 1.50 | 0.07 |
| 5B | 91.55 | 2.10 | 0.01 |
| 5B | 116.04 | 1.09 | 0.00 |
| 7B | 81.18 | 1.94 | 0.10 |
| 7B | 88.27 | 3.65 | 0.00 |
| 7B | 98.85 | 2.36 | 0.00 |

Supplementary Table 7: QTL mapping results, where trait is the number of recombination events across the whole genome.

| Subpopulation | AC-Barrie | Alsen | Baxter | Pastor | Volcani | Westonia | Xiaoyan | Yitpi |
|---|---|---|---|---|---|---|---|---|
| All | 0.19 | 0.12 | 0.11 | 0.07 | 0.18 | 0.07 | 0.15 | 0.11 |
| MP8RIL | 0.19 | 0.13 | 0.10 | 0.06 | 0.18 | 0.07 | 0.15 | 0.12 |
| AIC2RIL | 0.19 | 0.10 | 0.11 | 0.08 | 0.19 | 0.08 | 0.14 | 0.10 |
| AIC3RIL | 0.19 | 0.11 | 0.12 | 0.07 | 0.19 | 0.05 | 0.16 | 0.11 |

Supplementary Table 8: Genetic composition at position 155 cM on chromosome 2D.

| Subpopulation | AC-Barrie | Alsen | Baxter | Pastor | Volcani | Westonia | Xiaoyan | Yitpi |
|---|---|---|---|---|---|---|---|---|
| All | 0.05 | 0.17 | 0.14 | 0.14 | 0.12 | 0.12 | 0.12 | 0.14 |
| MP8RIL | 0.06 | 0.16 | 0.13 | 0.13 | 0.12 | 0.12 | 0.13 | 0.13 |
| AIC2RIL | 0.06 | 0.21 | 0.13 | 0.12 | 0.11 | 0.11 | 0.12 | 0.13 |
| AIC3RIL | 0.03 | 0.18 | 0.15 | 0.16 | 0.12 | 0.13 | 0.08 | 0.15 |

Supplementary Table 9: Genetic composition at position 136 cM on chromosome 4A.

| Subpopulation | AC-Barrie | Alsen | Baxter | Pastor | Volcani | Westonia | Xiaoyan | Yitpi |
|---|---|---|---|---|---|---|---|---|
| All | 0.09 | 0.22 | 0.17 | 0.11 | 0.16 | 0.08 | 0.09 | 0.09 |
| MP8RIL | 0.09 | 0.21 | 0.17 | 0.10 | 0.14 | 0.10 | 0.08 | 0.10 |
| AIC2RIL | 0.08 | 0.22 | 0.16 | 0.13 | 0.20 | 0.06 | 0.10 | 0.06 |
| AIC3RIL | 0.09 | 0.24 | 0.15 | 0.13 | 0.18 | 0.05 | 0.11 | 0.06 |

Supplementary Table 10: Genetic composition at position 72 cM on chromosome 7D.

| Subpopulation | AC-Barrie | Alsen | Baxter | Pastor | Volcani | Westonia | Xiaoyan | Yitpi |
|---|---|---|---|---|---|---|---|---|
| All | 0.15 | 0.12 | 0.10 | 0.09 | 0.14 | 0.07 | 0.17 | 0.15 |
| MP8RIL | 0.14 | 0.12 | 0.10 | 0.09 | 0.14 | 0.07 | 0.18 | 0.15 |
| AIC2RIL | 0.14 | 0.09 | 0.15 | 0.10 | 0.14 | 0.07 | 0.17 | 0.14 |
| AIC3RIL | 0.17 | 0.15 | 0.11 | 0.08 | 0.14 | 0.06 | 0.14 | 0.16 |

Supplementary Table 11: Genetic composition at position 238 cM on chromosome 7D.

| | AC-Barrie | Alsen | Baxter | Pastor | Volcani | Westonia | Xiaoyan | Yitpi | Total |
|---|---|---|---|---|---|---|---|---|---|
| AC-Barrie | 0.01 | 0.01 | 0.00 | 0.01 | 0.01 | 0.01 | 0.03 | 0.01 | 0.10 |
| Alsen | 0.01 | 0.01 | 0.00 | 0.01 | 0.01 | 0.01 | 0.02 | 0.01 | 0.08 |
| Baxter | 0.03 | 0.03 | 0.07 | 0.03 | 0.02 | 0.02 | 0.01 | 0.03 | 0.25 |
| Pastor | 0.01 | 0.01 | 0.00 | 0.01 | 0.01 | 0.02 | 0.03 | 0.01 | 0.12 |
| Volcani | 0.01 | 0.02 | 0.00 | 0.02 | 0.01 | 0.01 | 0.02 | 0.01 | 0.10 |
| Westonia | 0.01 | 0.01 | 0.00 | 0.01 | 0.01 | 0.01 | 0.02 | 0.01 | 0.09 |
| Xiaoyan | 0.02 | 0.02 | 0.00 | 0.02 | 0.02 | 0.02 | 0.04 | 0.02 | 0.16 |
| Yitpi | 0.01 | 0.01 | 0.00 | 0.01 | 0.01 | 0.01 | 0.03 | 0.01 | 0.10 |
| Total | 0.12 | 0.13 | 0.08 | 0.13 | 0.11 | 0.11 | 0.21 | 0.12 | |

Supplementary Table 12: Observed distribution of alleles, for the most significant 2B-2D interaction. Rows represent the genotype at position 445 cM on chromosome 2B. Columns represent the genotype at position 127 cM on chromosome 2D.

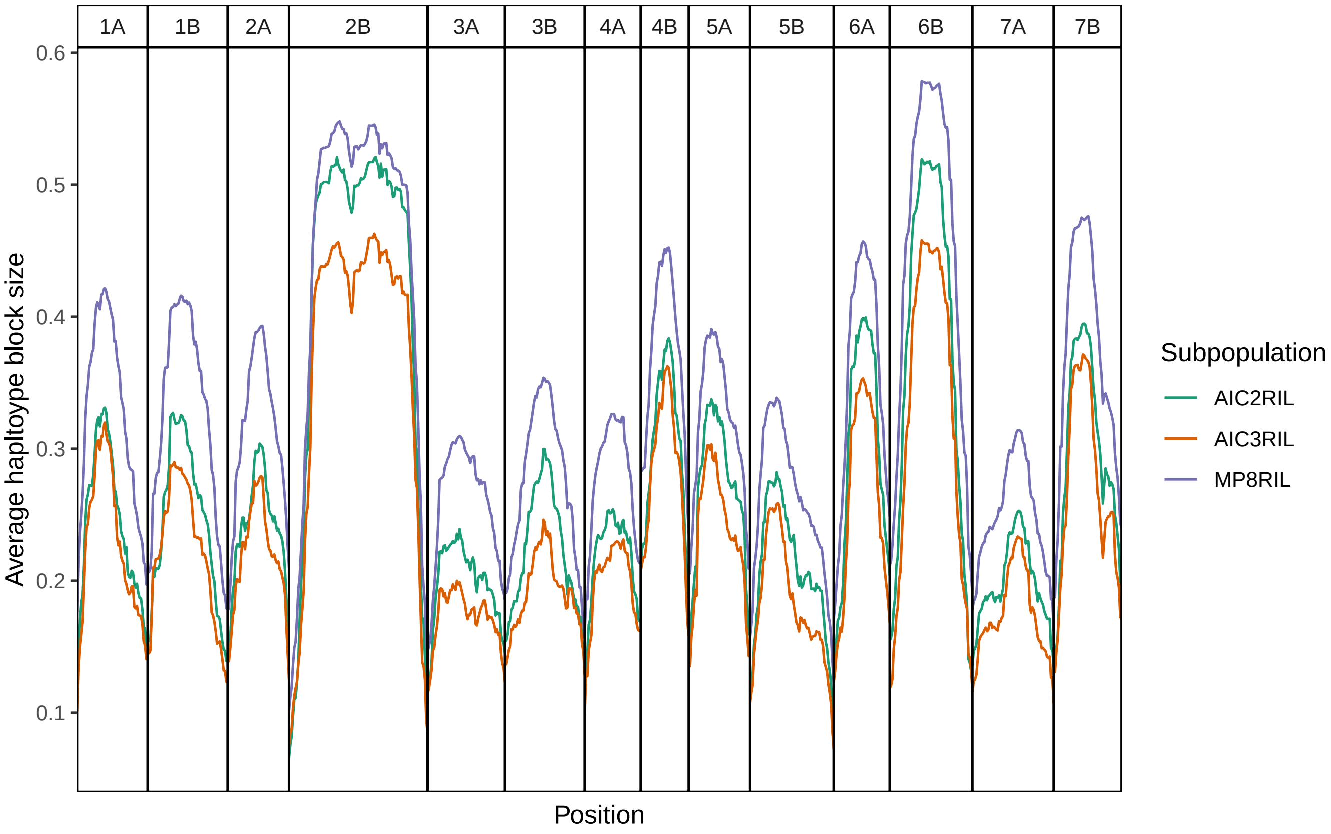


Supplementary Figure 10: Average haplotype block sizes for the A and B genomes, as a proportion of the total chromosome length.

| | AC-Barrie | Alsen | Baxter | Pastor | Volcani | Westonia | Xiaoyan | Yitpi | Total |
|---|---|---|---|---|---|---|---|---|---|
| AC-Barrie | 0.01 | 0.01 | 0.02 | 0.01 | 0.01 | 0.02 | 0.01 | 0.01 | 0.09 |
| Alsen | 0.01 | 0.01 | 0.02 | 0.01 | 0.01 | 0.02 | 0.01 | 0.01 | 0.10 |
| Baxter | 0.02 | 0.01 | 0.05 | 0.01 | 0.02 | 0.05 | 0.01 | 0.05 | 0.23 |
| Pastor | 0.01 | 0.01 | 0.02 | 0.01 | 0.01 | 0.03 | 0.01 | 0.02 | 0.12 |
| Volcani | 0.01 | 0.02 | 0.02 | 0.02 | 0.01 | 0.02 | 0.01 | 0.02 | 0.13 |
| Westonia | 0.01 | 0.01 | 0.02 | 0.01 | 0.01 | 0.02 | 0.01 | 0.02 | 0.09 |
| Xiaoyan | 0.01 | 0.01 | 0.02 | 0.02 | 0.01 | 0.02 | 0.01 | 0.03 | 0.14 |
| Yitpi | 0.01 | 0.01 | 0.02 | 0.02 | 0.01 | 0.01 | 0.01 | 0.01 | 0.10 |
| Total | 0.09 | 0.10 | 0.19 | 0.09 | 0.09 | 0.19 | 0.07 | 0.18 | |

Supplementary Table 13: Observed distribution of alleles, for the most significant 2B-6B interaction. Rows represent the genotype at position 462 cM on chromosome 2B. Columns represent the genotype at position 306 cM on chromosome 6B.

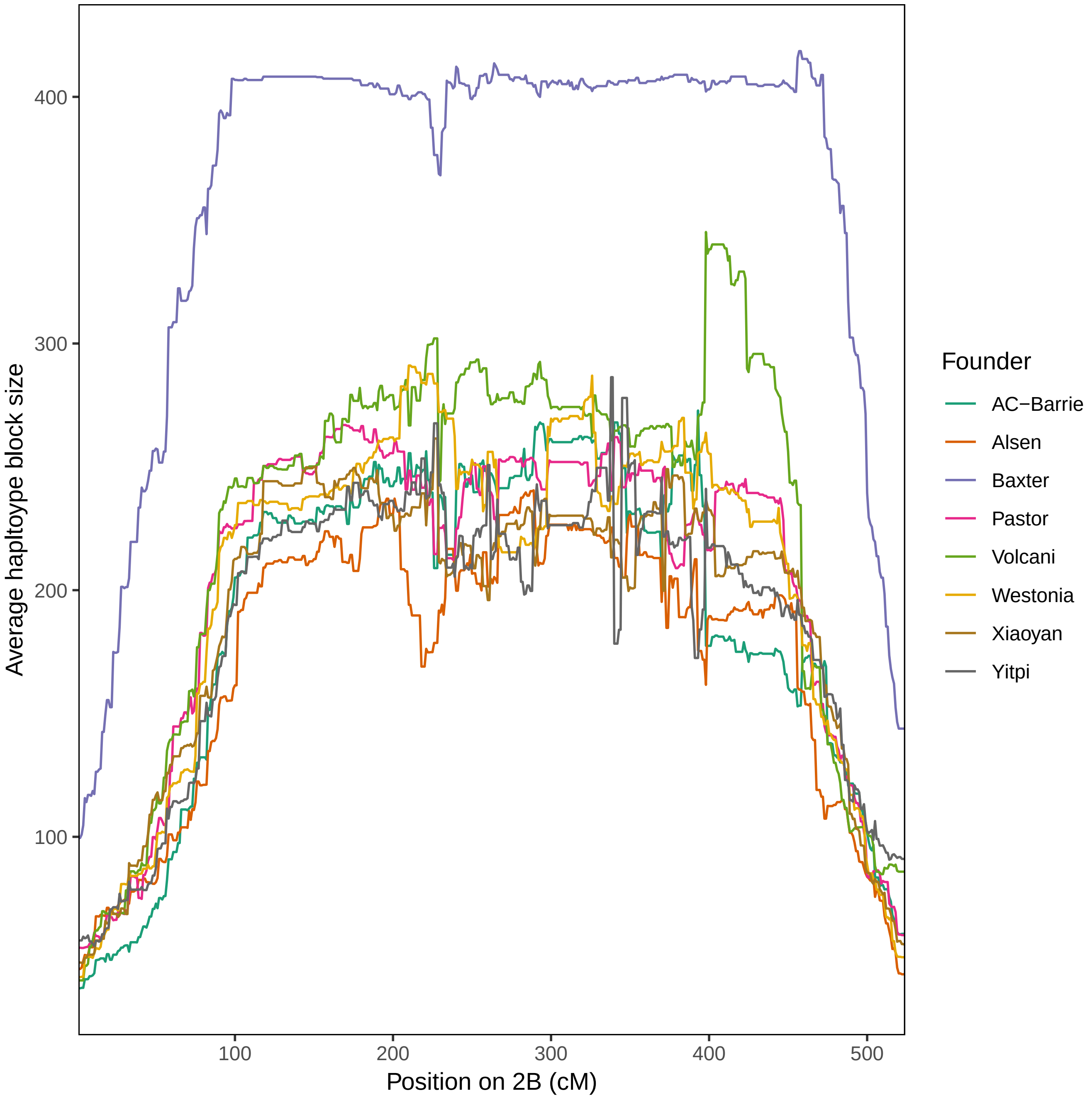


Supplementary Figure 11: Average haplotype block sizes for each founder, at different positions on chromosome 2B.

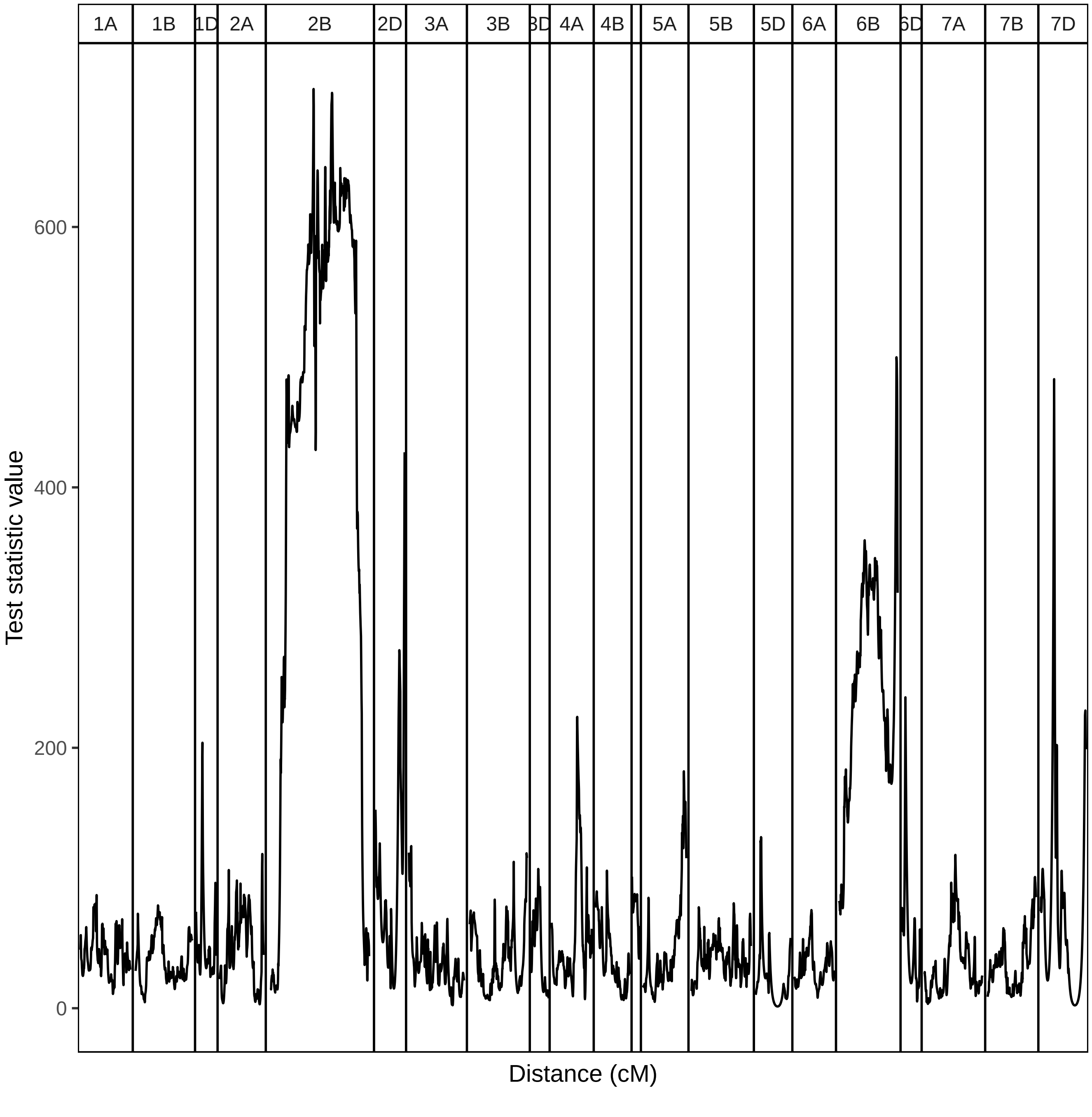


Supplementary Figure 12: Chi-squared test statistic values, for tests of segregation distortion. Chromosome 4D is not labelled due to the width of the panel.

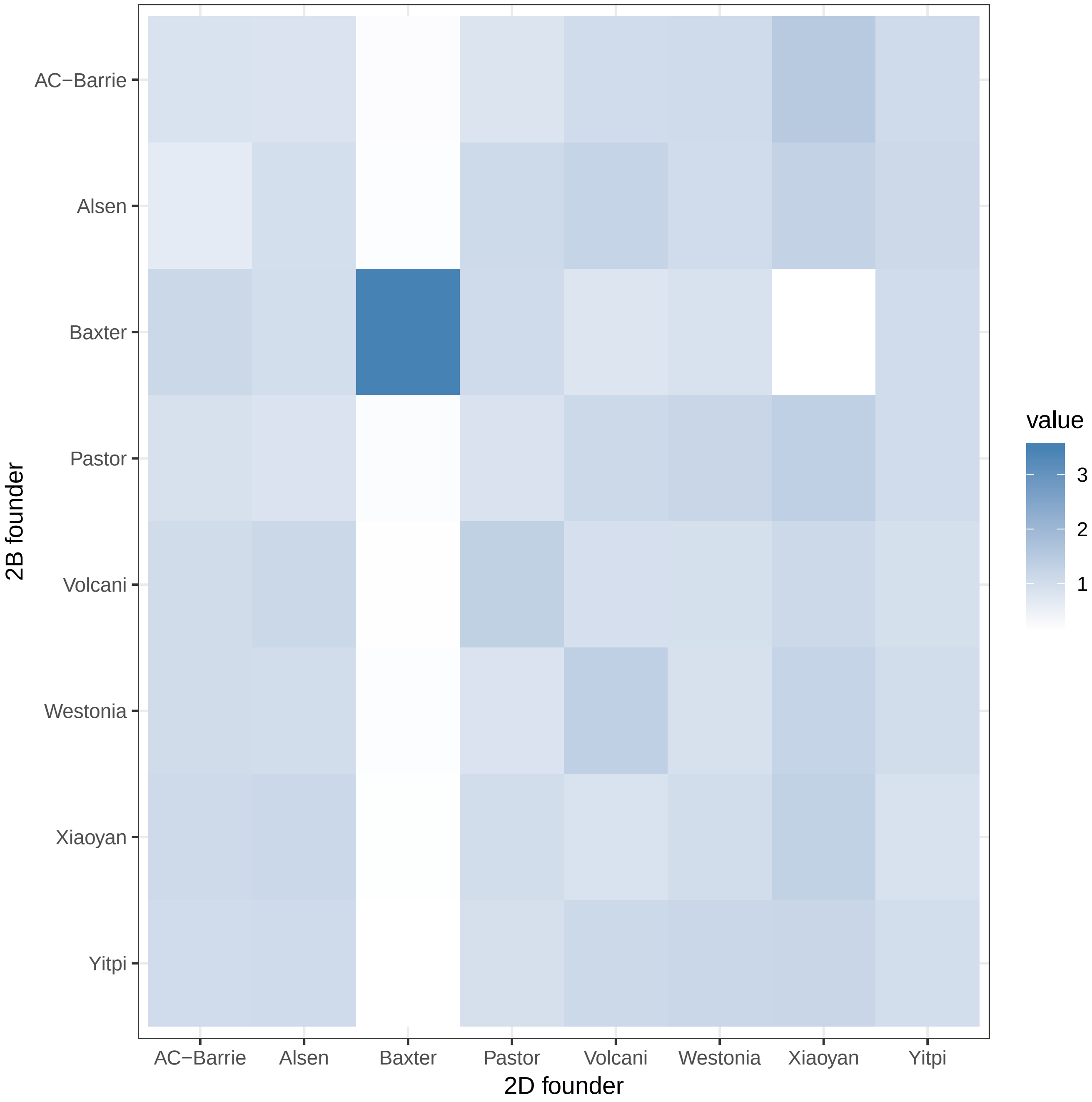


Supplementary Figure 13: Visualisation of the 2B-2D interaction in Table 12. Colour represents the observed frequency as a multiple of the expected frequency under independence. Dark blue represents combinations present much more frequently than expected.

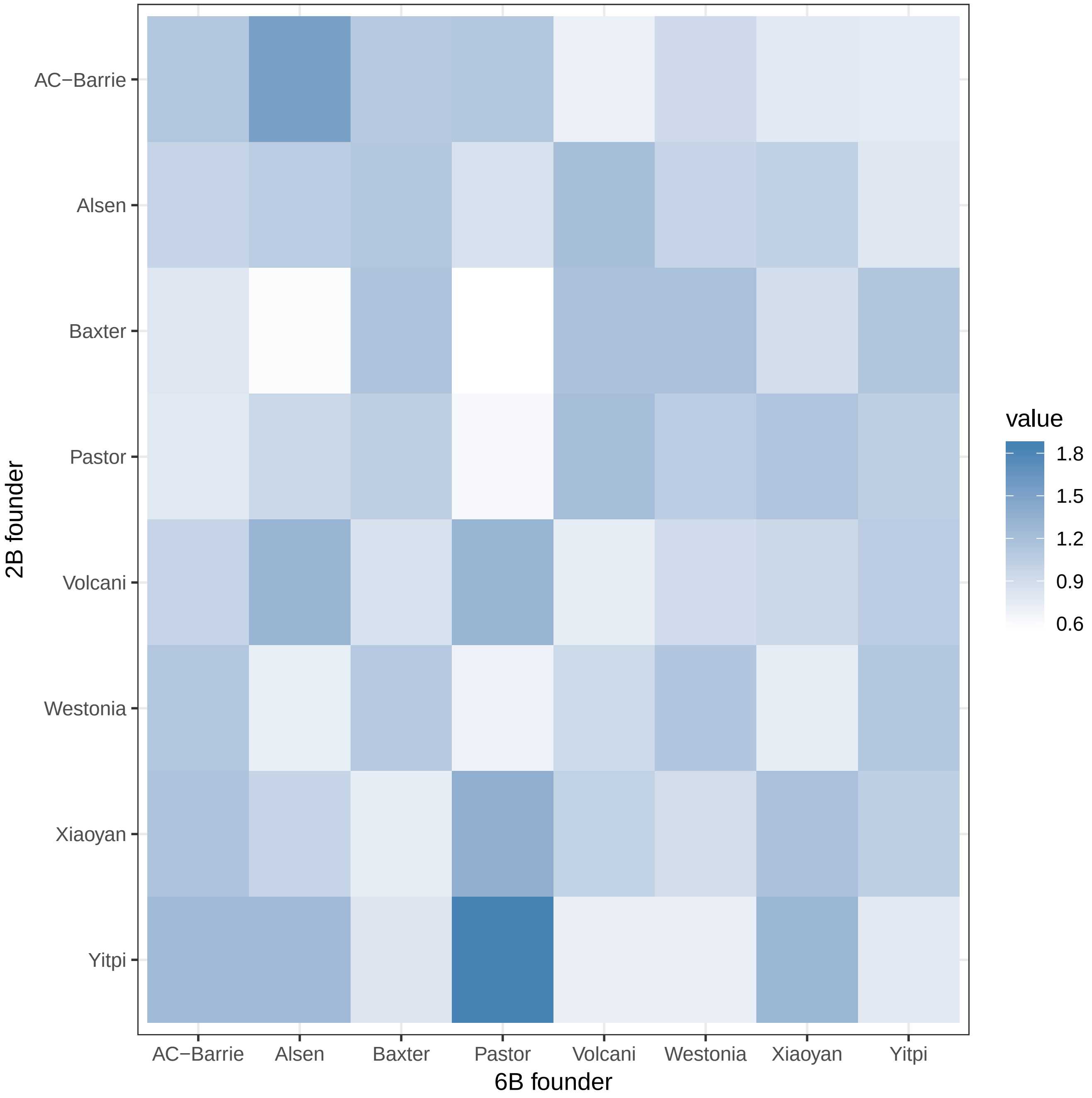


Supplementary Figure 14: Visualisation of the 2B-6B interaction in Table 13. Colour represents the observed frequency as a multiple of the expected frequency under independence. Dark blue represents combinations present much more frequently than expected.

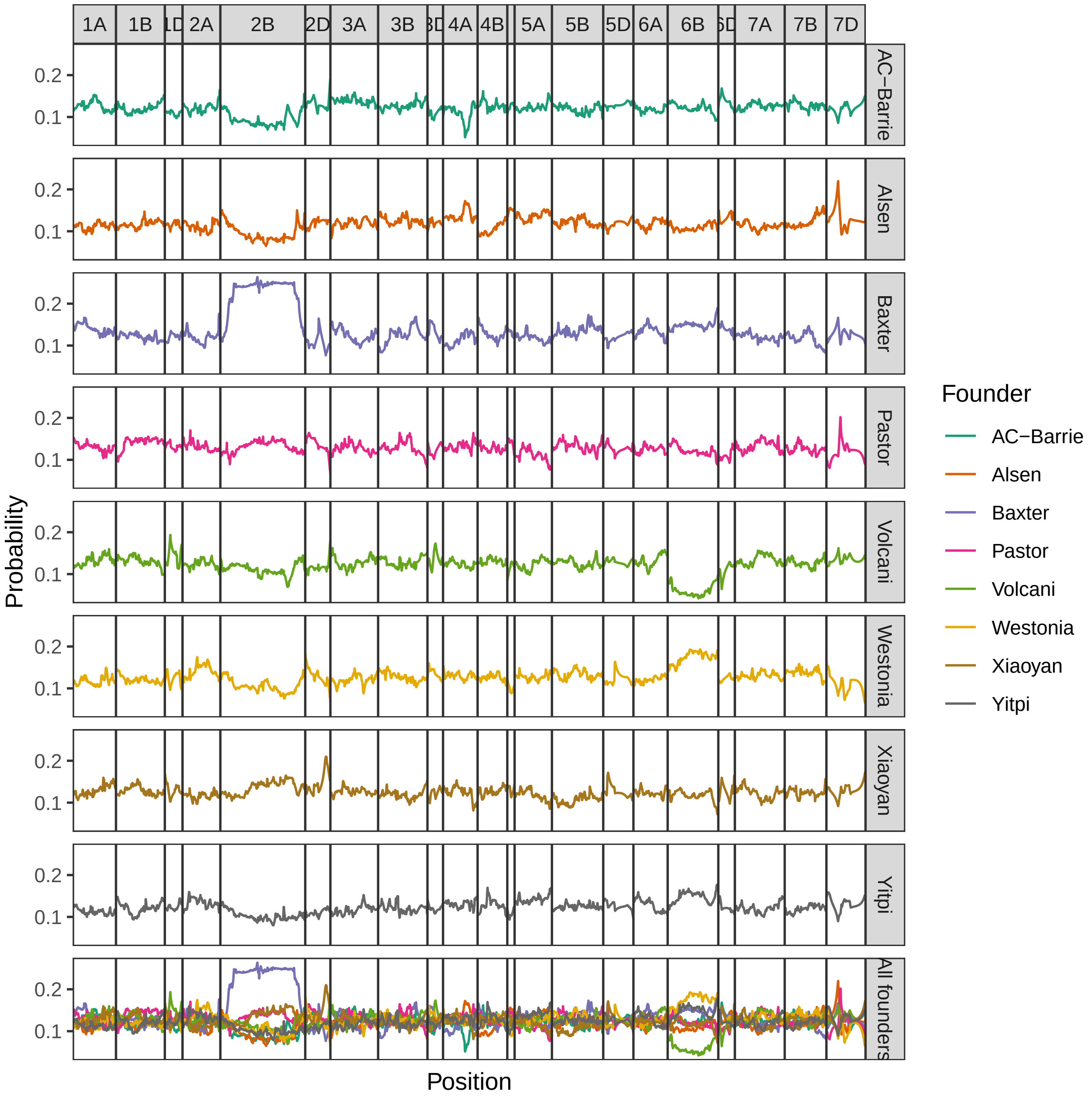


Supplementary Figure 15: Average genetic composition across the entire population

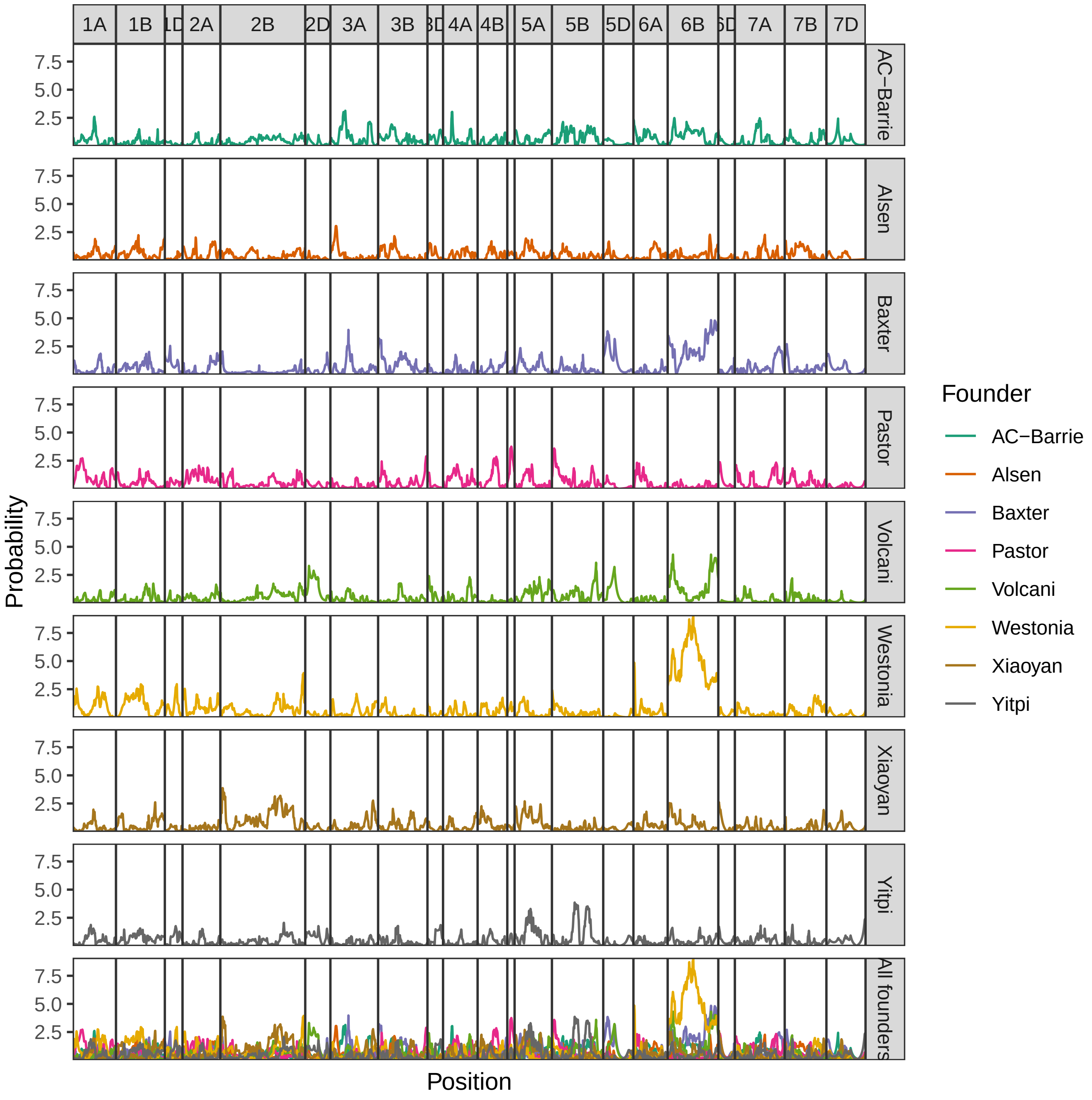


Supplementary Figure 16: Chi-squared statistics for sex-specific segregation distortion.

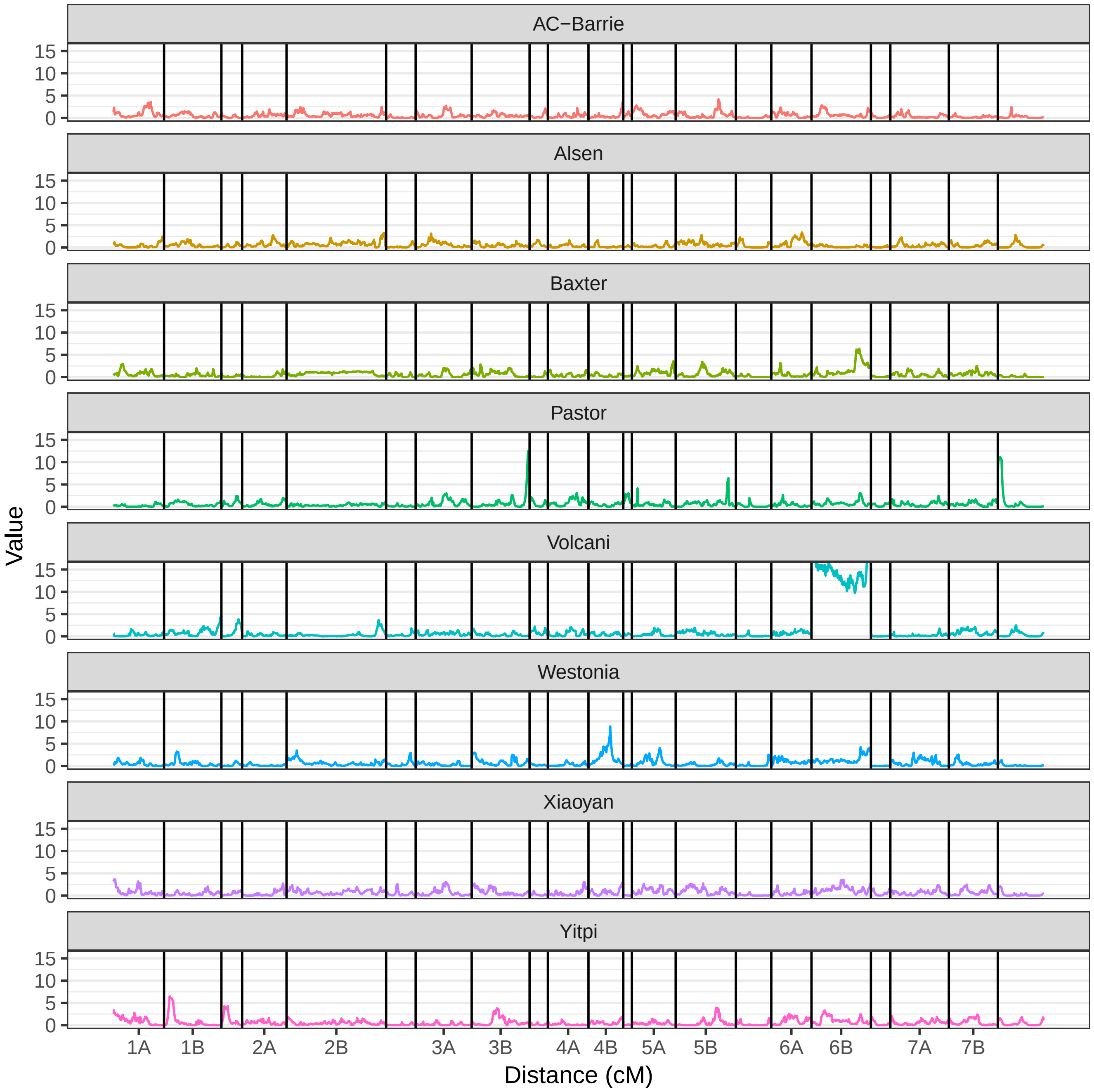


Supplementary Figure 17: Chi-squared statistics for segregation distortion related to the crossing of founders in the first generation.

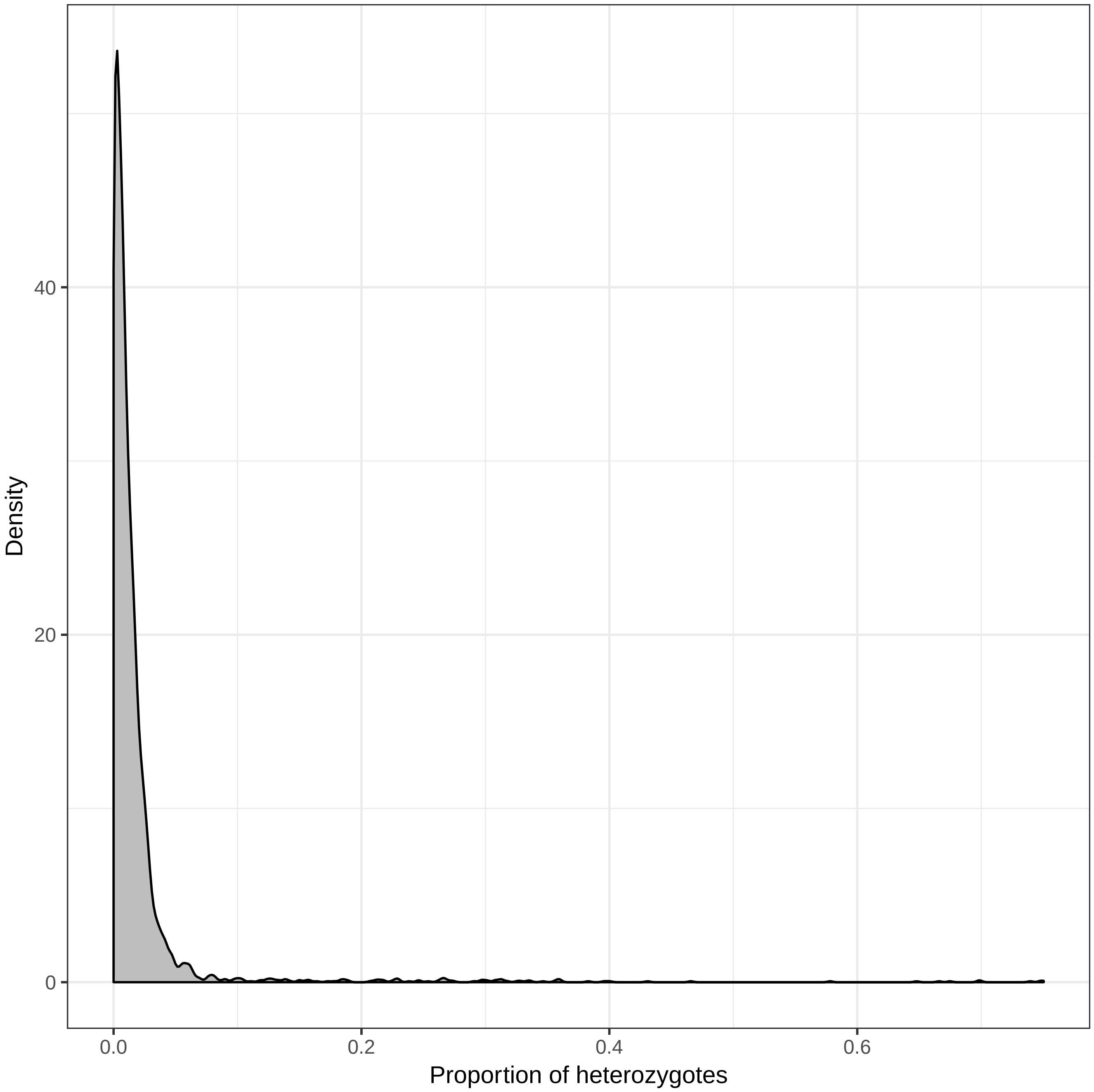


Supplementary Figure 18: Distribution of the proportion of residual heterozygosity, for all three subpopulations.

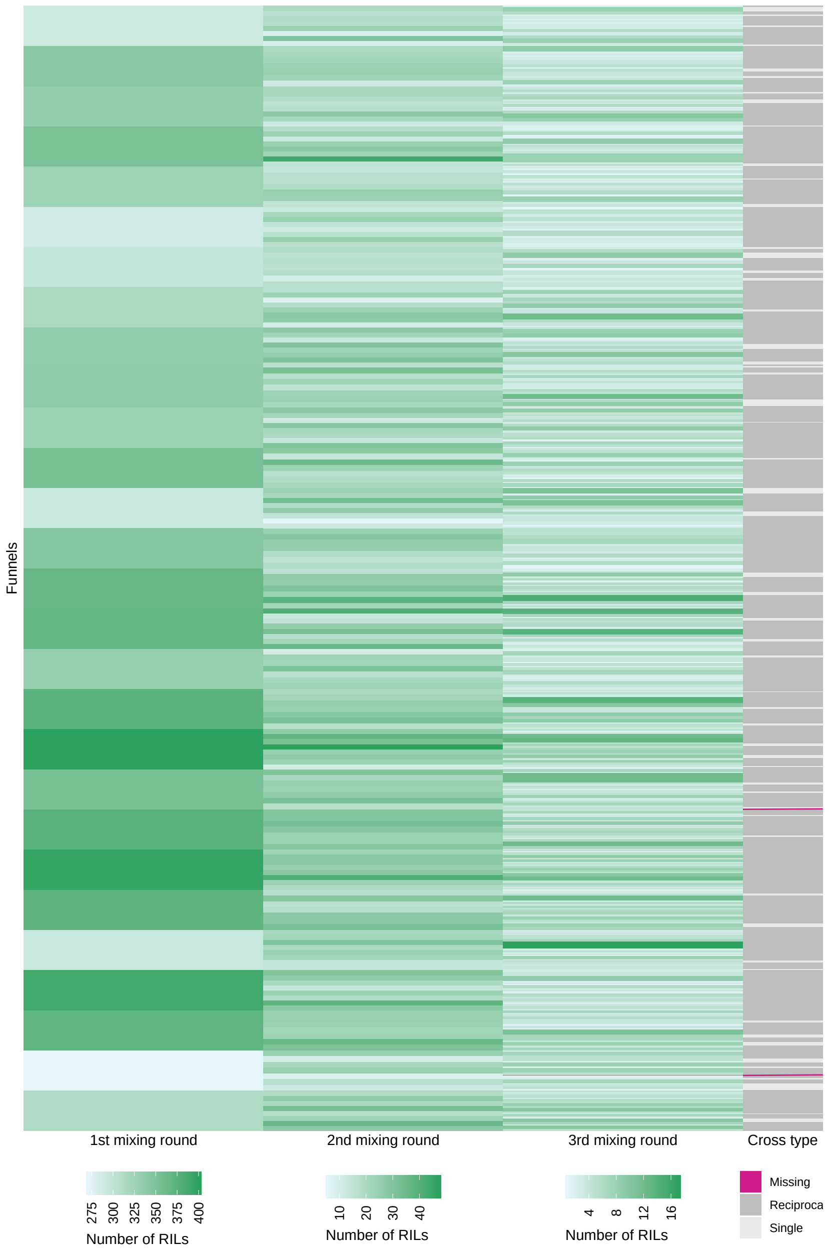


Supplementary Figure 19: Heatmap representation of funnels in final population, according to the maternal parent in each round. Each rectangle represents a single cross. The rectangles are coloured according to how many MP8RILs were derived from that cross in the final population. The rectangles are arranged in funnels so that crosses in each round of mixing are positioned to the right of the maternal cross from the previous round. The two missing crosses in the third round of mixing are highlighted in red. At the far right, crosses in the third round that were performed in both directions (reciprocal crosses), are shown by a grey line.

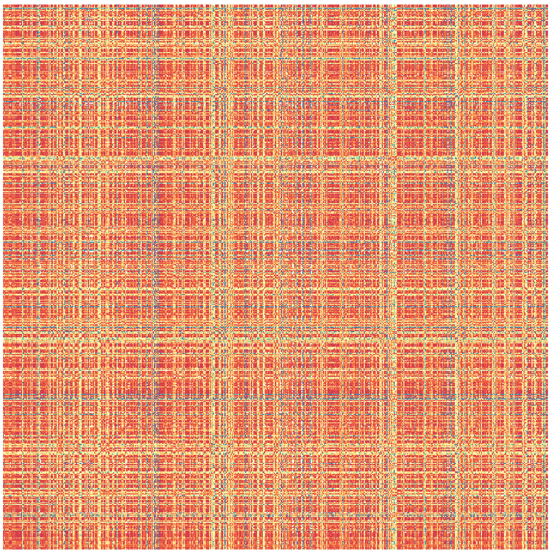

(a) Before ordering. Markers are in a random order.

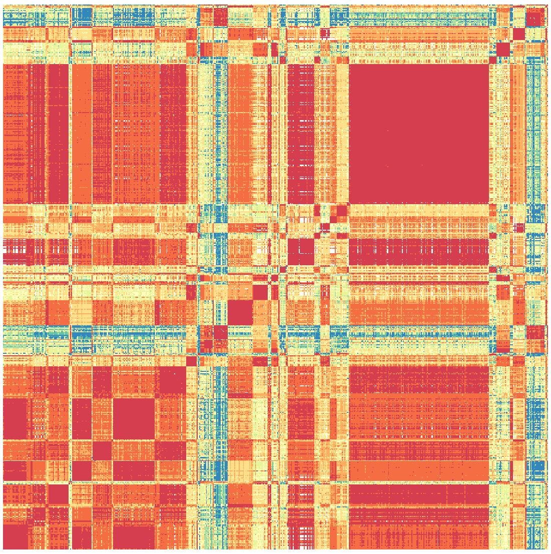

(b) After ordering the markers into 50 blocks. Markers within blocks are unordered.

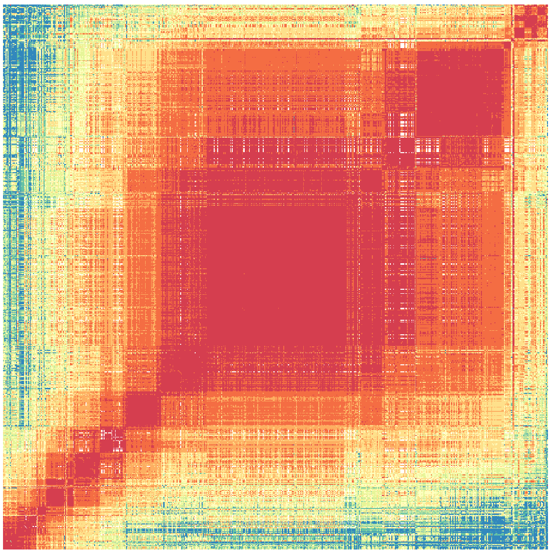

(c) After ordering the markers into 50 blocks, and then ordering the 50 blocks. Markers within blocks are still unordered.

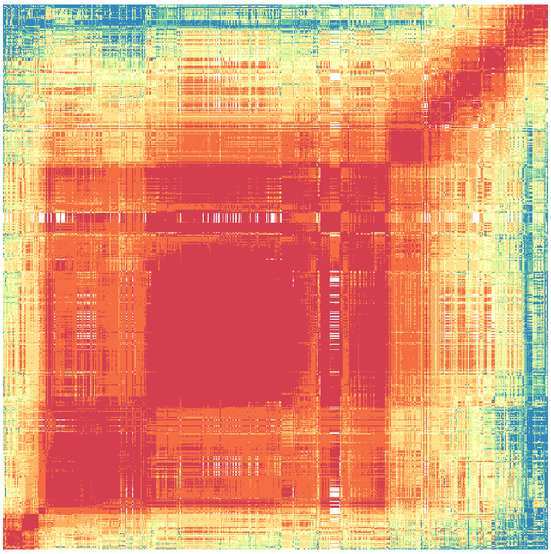

(d) A close to optimal ordering for this particular set of markers.

Supplementary Figure 20: Rows and columns represent genetic markers on this chromosome, and values in the image represent estimated recombination fraction between a pair of markers. The ordering of the markers is different in all four subfigures. Red represents small recombination fractions, while blue represents large values. (a) The ordering of markers is initially random. (b) The markers are grouped into 50 large blocks. (c) These blocks are then ordered. (d) A close to optimal ordering.
